# Optimized single molecule fluorescence sheds light on elusive enzymatic mechanisms

**DOI:** 10.1101/2020.01.16.908947

**Authors:** Marko Usaj, Luisa Moretto, Venukumar Vemula, Aseem Salhotra, Alf Månsson

## Abstract

Single molecule enzymology using fluorescent substrate requires truly minimal amounts of proteins. This is highly beneficial when the protein source is either advanced expression systems or samples from humans/animals with ethical and economic implications. Further benefits of single molecule analysis is the potential to reveal phenomena hidden in ensemble studies. However, dye photophysics and fluorescent contaminants complicate interpretation of the single molecule data. We here corroborate the importance of such complexities using fluorescent Alexa647 ATP to study ATP turnover by myosin and actomyosin. We further show that the complexities are largely eliminated by aggressive surface cleaning and use of a range of triple state quenchers and redox agents with minor effects on actin-myosin function. Using optimized assay conditions, we then show that the distributions of ATP binding dwell times on myosin are best described by the sum of 2 to 3 exponential processes. This applies in the presence and absence of actin and in the presence and absence of the drug para-aminoblebbistatin. Two of the processes are attributable to ATP turnover by myosin and actomyosin, respectively. A remaining process with rate constant in the range 0.2-0.5 s^-1^ is consistent with non-specific ATP binding to myosin and bioinformatics modelling suggests that such binding may be important for accelerated ATP transport to the active site. Finally, we report studies of the actin-activated myosin ATP turnover under conditions with no sliding between actin and myosin, as in isometrically contracting muscle, revealing heterogeneity in the ATP turnover kinetics between different molecules.

## Introduction

Extensive miniaturization by single molecule methods reduces the need of biological material in biochemical assays with economic, ethical and scientific implications. The first fluorescence based single biomolecule assay (1) used total internal reflection fluorescence (TIRF) microscopy to study the turnover of a fluorescent ATP analog by the mechano-enzyme myosin II. This molecular motor, underlying muscle contraction and cell motility, is an emerging drug target in severe diseases such as cancer (2, 3), heart failure (4, 5) and hypertrophic cardiomyopathy (6) making single molecule ATPase assays of growing interest. However, recent observations of additional exponential processes (7–9) and ~5-50-fold faster rate constants in single molecules (7–9) compared to ensemble studies (10, 11) are problematic in this context. Such inconsistencies also hamper the potential to derive rich and reliable mechanistic information from single molecule fluorescence data (12–16). Notably, the fast processes, found recently (7–9), have a counterpart in early reports from the 1990s. At the time, however, the faster events were attributed to intrinsic characteristics of expressed constructs (i.e. lack of the light-chain binding neck domain) (15) or to assay limitations (i.e. rebinding of ADP or presence of contaminating particles). The fast events were therefore excluded from the analysis as they were not believed to reflect enzymatic mechanisms (13). In contrast, recent work (7) associated the fast processes with myosin conformers having different catalytic activity. Clearly, both for the general usefulness of single molecule data in applications e.g. drug screening and for deriving reliable mechanistic information, it is essential with unequivocal understanding of the fast processes. Naturally, if they just reflect artefacts they should be excluded from the analysis (13) if not, they may convey important mechanistic information per se (7).

We here aim to identify the basis for the fast events mentioned above, and provide a solid basis for reliable single molecule assays. Our results corroborate a hypothesis that the faster rate constants reflect a combination of dye photophysics events (i.e. photoblinking) (17, 18) and surface contamination with unidentified fluorescent objects. However, interestingly, despite optimizations to remove these confounding effects we still observe one unexplained process in addition to that consistent with the ATP turnover rate in solution (7–9). Experimental tests and bioinformatics modelling attribute this unexplained process to non-specific binding of ATP with a role in accelerating ATP-transport to the active site. In addition to the studies of myosin, our assay optimizations allow unique insights into the >10-fold faster actomyosin ATPase, e.g. revealing heterogeneity between individual molecules. This heterogeneity is most likely not due to a dynamic shift between different myosin conformers. As further expanded on below, it is instead accounted for within an existing theoretical framework as being due to strain dependence of the detachment rate of myosin from actin.

## Results and Discussion

### Single molecule basal myosin ATPase

We used objective based TIRF microscopy (Fig. S1) of the fast skeletal muscle myosin II motor fragments heavy meromyosin (HMM) and subfragment 1 (S1) adsorbed to silanized glass surfaces (Fig. 1a-b). The fluorescent ATP analogue Alexa647-MgATP (“Alexa-ATP”) that we use, works virtually identically as non-fluorescent MgATP with respect to ATP-turnover kinetics and powering of myosin driven actin filament sliding velocity (19). Furthermore, in contrast to conventionally used Cy3 based ATP analogues (1, 7, 8, 20), turnover kinetics are similar between different Alexa-ATP isomers (19). We performed initial studies in standard degassed in vitro motility assay (IVMA) solution (cf. (7)) containing GOC (glucose, glucose oxidase and catalase) and DTT as oxygen scavenger and reducing agent, respectively. As in previous work (7), fluorescence dwell-time events (Fig. 1c) were collected from surface “hotspots” attributed to myosin motor domains and the resulting cumulative dwell-time distributions were fitted by double exponential functions (Fig. 1e-f). The rate constants (~5 s^-1^; ~60 % and ~0.5 s^-1^; ~40 %) were appreciably faster than kcat in solution (0.05-0.1 s^-1^), corroborating the inconsistencies between single molecules and ensemble data previously demonstrated using Cy3-ATP (7) and similar assay solution (IVMA solution).

**Figure 1.**
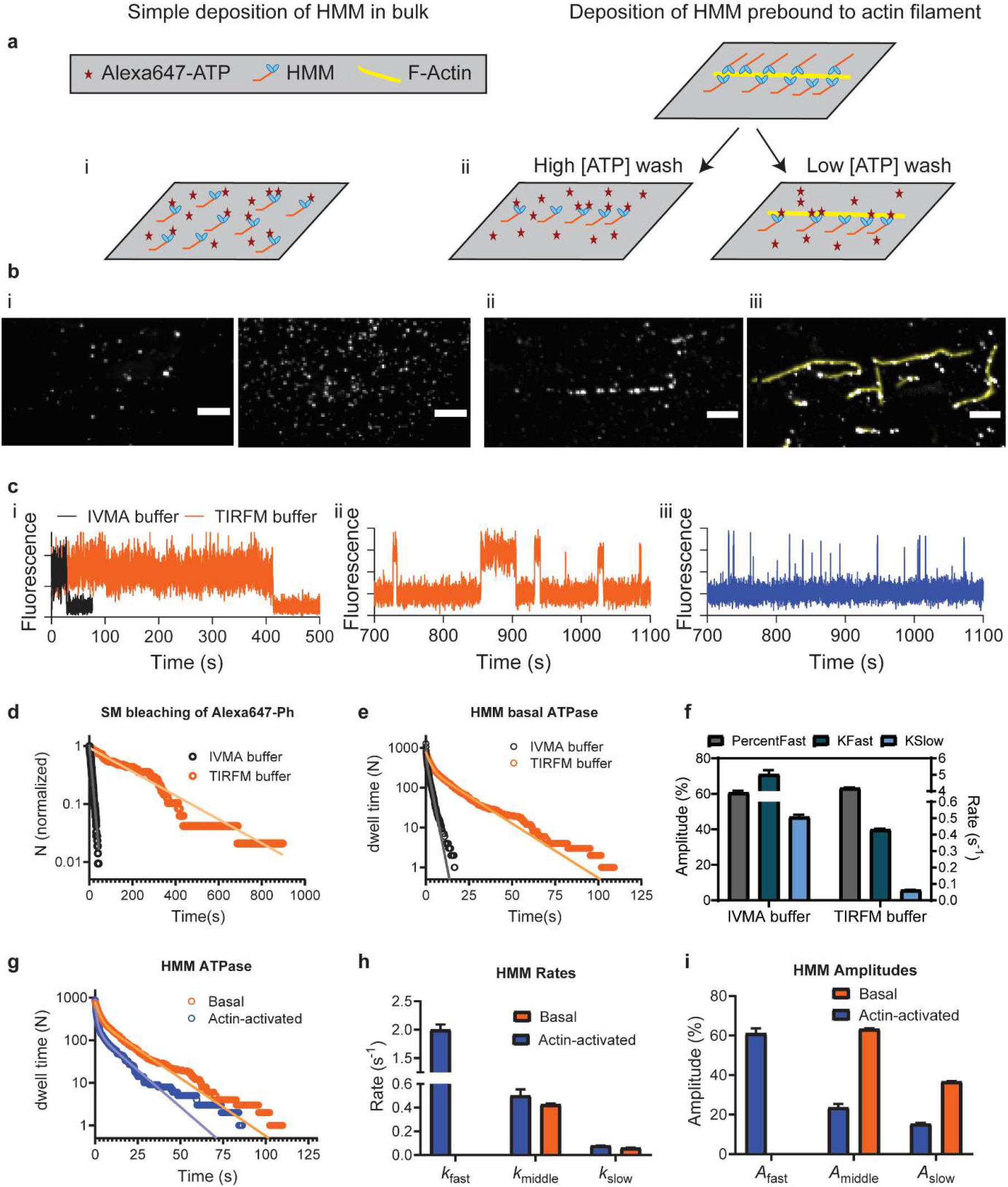
Single molecule ATPase. **a, Two principles for surface immobilization of myosin motor fragments.** b, Time-averaged Alexa-ATP fluorescence projections of 15 min videos (50 ms exposure time/frame). i, Simple HMM deposition at 34.3 pM (left) and 343 pM (right). ii, Optimized HMM deposition following high [ATP] wash. Only spots defined by previous position of F-actin filament were included in analysis. iii, Image of optimized actomyosin deposition followed by low [ATP] wash showing co-localization of Alexa-ATP fluorescence (grey spots) and F-actin filaments (yellow). Only spots co-localizing with Factin filaments were included in analysis. Bars, 5 μm. c, Representative time traces of i, single molecule Alexa-Ph bleaching, ii, HMM basal ATPase and iii, actin activated HMM ATPase. Bleaching was observed either using standard in vitro motility assay (IVMA) buffer or optimized TIRF microscopy buffer (TIRFM) as described in the text whereas ATPase traces were from experiments using TIRFM buffer. d, Cumulative frequency distributions of single molecule Alexa-Ph until bleach or first blinking event. The distributions are fitted by single exponential functions (solid lines). e, Cumulative frequency distribution of Alexa-nucleotide dwell-time events on HMM surface hot-spots (IVMA buffer-simple deposition of HMM, TIRFM buffer-optimized deposition of HMM) were fitted with double exponential functions (solid lines). IVMA data from 68 HMM molecules, N_dwell_ = 1273, TIRFM data from 45 HMM molecules, N_dwell_ = 785. f, Amplitudes and rate constants from the fitting of data in e. Note appreciably lower rate constant values in TIRFM compared to IVMA buffer. g, Cumulative frequency distributions of Alexa-nucleotide dwell-time events comparing HMM basal and actin-activated ATPase activity. Actin-myosin data fitted with triple exponential function whereas basal ATPase is fitted by double exponential function. h, Rate constants obtained from fittings to data in f. i, Amplitudes from fitting the data in f. Error estimates refer to 95 % confidence intervals derived in the regression analysis. Temperature: 23 °C.

### Single molecule assay optimizations

As first optimization steps, we strived to remove unidentified fluorescent objects on motor adsorbing surfaces by extensive surface cleaning (21) and refined selection and processing of the bovine serum albumin (BSA) (22) (Fig. S2) used for surface blocking. To characterize dye photophysics, isolated from turnover events, we next studied Alexa647-phalloidin (“Alexa-Ph”) sparsely bound to actin. By sequential addition of redox-agents and triple state quenchers (Cyclooctatetraene, 4-Nitrobenzyl alcohol and optimized mixtures of trolox and trolox-hydroquinone) (18) (Fig. S3), the rate constant for single molecule photobleaching/blinking was progressively reduced (Fig. 1d) more than 10-fold down to 0.004 ±0.00003 s^-1^ (mean ± 95 % CI; Fig. S3). Importantly, Alexa-ATP locked into the nucleotide pocket by Vanadate (23, 24), exhibited similar photobleaching rate constant as Alexa-Ph, suggesting minor effects of the dye microenvironment. However, the intensity traces (Fig. S4) with Alexa-ATP were noisier than with Alexa-Ph. We attribute this effect to thermal fluctuations (23) as it was reduced by denser packing of HMM on the surface or increased solution viscosity using methylcellulose. To be fully confident that observed hotspots represent HMM molecules, we adsorbed HMM via actin filaments (Fig. 1a) and limited the analysis to positions defined by the filaments (Fig. 1b). We also took extra care to exclude any remaining effects of non-specific surface/BSA binding of Alexa-ATP or the Alexa moiety itself (Fig. S5). Because our optimized TIRF assay buffer differs from the standard IVMA assay buffer, we performed IVMA tests in order to examine any adverse effect of the new additives. The addition of all redox-agents and triple state quenchers together had mild inhibitory effect on actomyosin function (Fig. 2) with a small reduction in actin filament velocity when the in vitro motility assay was performed in TIRF buffer instead of standard IVMA buffer. Reduced velocity (Fig. 2a) can be attributed to the effect of DMSO (2 %), used as a vehicle for COT and NBA and known as reversible inhibitor of actomyosin function (25–29). Importantly, however, the fraction of motile filaments (FMFs) was not affected (Fig. 2b).

**Figure 2.**
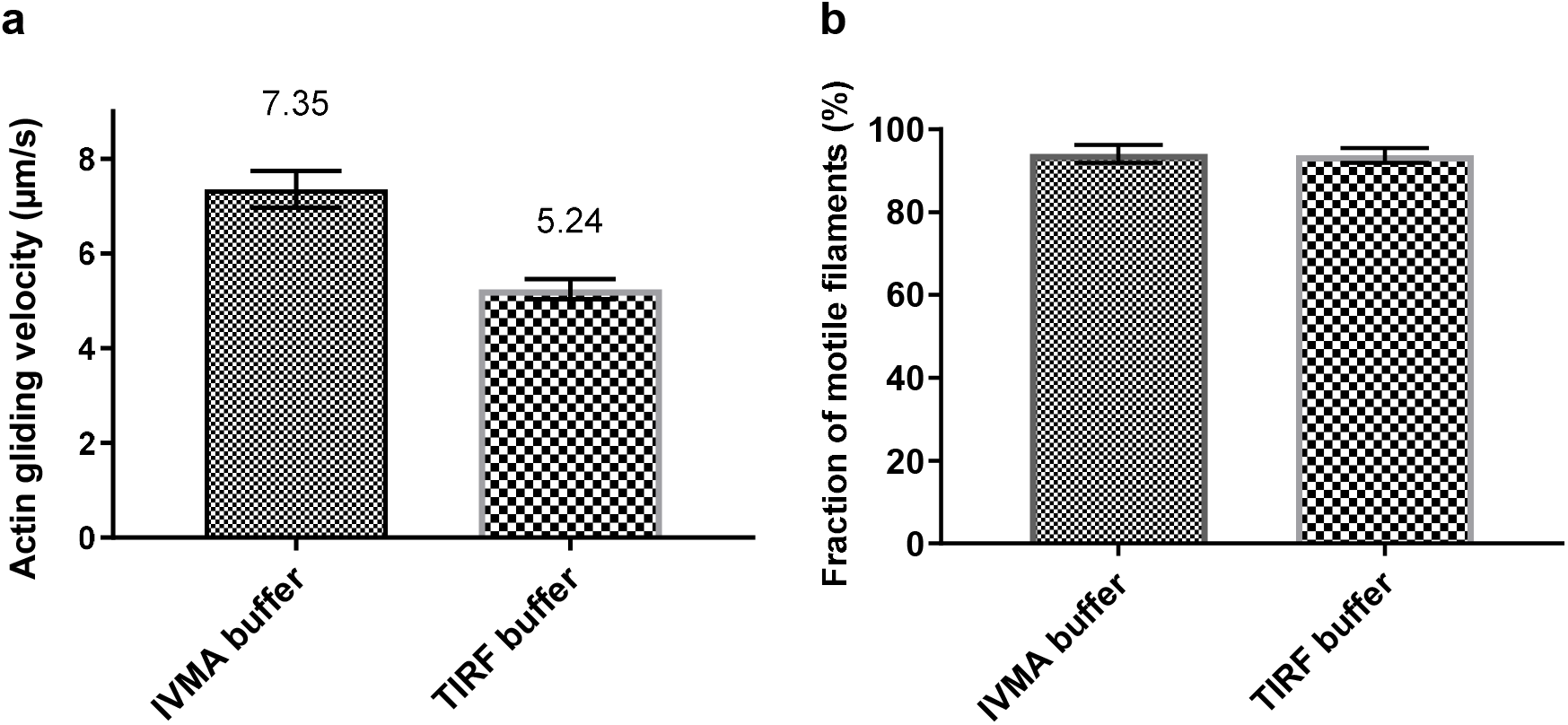
Analysis of motility in standard IVMA buffer and optimized TIRF buffer. **a,** Actin gliding velocity produced by HMM adsorbed to TMCS derivatized glass surfaces. In each individual experiment, 15 actin filaments were analyzed. **b,** Fraction of motile filaments (FMFs) from the same experiments analyzed from total 1004 filaments (IVMA buffer) and 925 filaments (TIRF buffer), respectively. Data are given as mean ± 95% confidence interval. Temperature 24-26 °C.

### Single molecule myosin basal and actin-activated ATPase under optimized conditions

Also after the methodological optimizations, dwell-time distributions using myosin motor fragments without actin (Fig. 1e-f, Fig. S6) were better fitted by a double than a single exponential function. However, the rate constants were reduced ~10-fold compared to the situation without optimizations (Fig. 1e-h). Particularly, the fastest phase (rate constant >2 s^-1^), ubiquitous and dominant in previous work (>70 %; (7)), disappeared and was not reinstated by triple exponential fitting (Fig. 2b, Fig. S7) or other changes in the fitting procedure (Fig. S7; Table S1). Instead, a slow phase emerged, consistent with basal MgATP turnover (10, 19, 30). The origin of the remaining unexplained phase (0.2-0.5 s^-1^; 40-70 % amplitude with HMM and S1; Figs. 1, S7) is considered below.

The optimizations allowed reliable studies also of the actin-activated myosin ATPase (Fig. 1b-c, g-i). The very low Alexa-ATP concentration (≤ 10 nM) gives “isometric” conditions with undetectable sliding of actin vs myosin. The triple exponential dwell-time distribution (Fig. 1g-i; S8) for actin-activated ATP turnover exhibited a fast rate constant of 2-3 s^-1^ (61-74 %), an intermediate rate constant of 0.4-0.5 s^-1^ (22-24 %) and a slow rate constant of 0.05-0.08 s^-1^ (4-15 %). Here, we attribute the fastest and slowest phase to actin-activated and basal myosin ATP turnover, respectively whereas the intermediate phase corresponds to the unexplained phase mentioned above. Notably, the actin-activated ATP turnover rate is slower than 10-17 s^-1^ for isometrically contracting muscle fibres ((31) and references therein) but quite similar to the rate (4 s^-1^) of the slow phase of isometric relaxation upon removal of calcium from a muscle cell (32). Because detachment from actin of high-force myosin heads dominates the latter process, it is reasonable to presume that our single molecule data primarily report turnover of high-force heads. A possible basis for faster average ATP turnover rate in muscle may be that all sarcomeres are not isometric but that some shorten against weaker, elongating, sarcomeres (33). However, distributions derived at single, individual actin-myosin interaction points (Fig. 3) demonstrate an appreciable heterogeneity in rate constants with some of the fast rate constants even approaching actin-activated ATP turnover rate for isometrically contracting muscle fibres. This heterogeneity is understandable on basis of strain dependent actomyosin detachment kinetics consistent with the detachment rate limitation of the ATP turnover by actomyosin under isometric conditions. A variability in strain between different actomyosin cross-bridges is expected to exist because the distance between the point of surface adsorption of the HMM molecule and the nearest myosin binding site on the actin filament varies.

**Figure 3.**
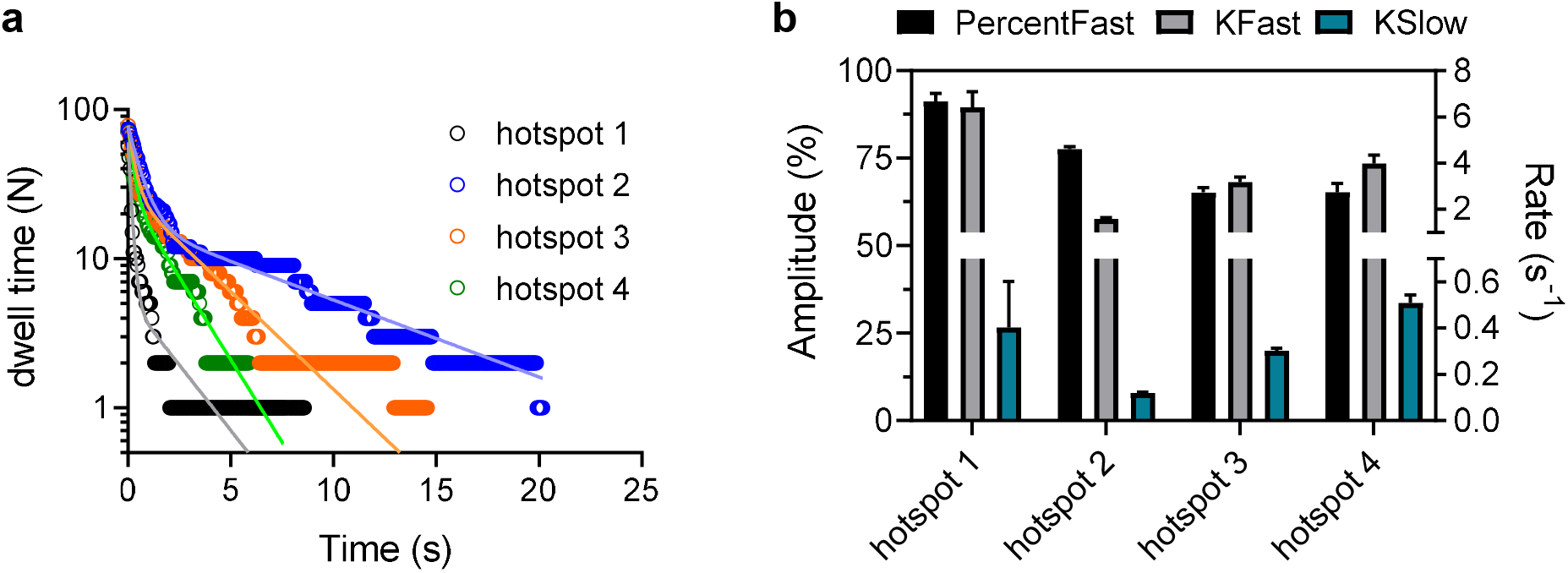
Cumulative dwell time distributions recorded from different individual actomyosin hotspots to illustrate the heterogeneity in time constants. **a,** Cumulative frequency distributions of Alexa-nucleotide dwell-time events comparing actin-activated ATPase activity from four different hotspots. Data are fitted with double exponential function. **b,** Amplitudes and rate constants obtained from fittings to data in a. Note that from the individual actomyosin hotspots except hotspot 2 only two processes attributed to actin-activated ATPase and “unexplained” phase can be resolved while basal myosin ATPase remains obscure. For hotspot 2 it seems likely that the binding site of the HMM molecule is unfavorable for actin attachment, explaining a high ratio of basal to actomyosin ATPase. Error estimates refer to 95 % confidence intervals derived in the regression analysis. Temperature: 20 °C.

Furthermore, strikingly, as discussed below, dwell-time records (Fig. 1c) suggest (from average waiting times between events at 10 nM Alexa-ATP) that the second order rate constant (K1k2) for Alexa-ATP binding to myosin increases from (2.80 ± 0.02) 10^6^ M^-1^s^-1^ (similar to solution data (34)) in the absence of actin to (4.74 ± 0.01) 10^6^ M^-1^s^-1^ in its presence. Importantly, our reliable analysis of the actomyosin MgATP turnover depends heavily on the methodological optimizations because the fast exponential processes previously found with myosin alone would otherwise have disturbed the analysis.

### Origin of unexplained phase in the Alexa-ATP dwell time distributions

One possible interpretation of the double-exponential dwell time distributions with myosin alone (Fig. 1) is that it is due to two myosin conformers (7) but with different kinetic properties than suggested in (7). Interestingly, however, our observation of non-specific binding of Alexa-ATP to BSA (Fig. S2; see also (35, 36)) suggested an alternative origin of the unexplained phase, namely non-specific binding of Alexa-ATP to myosin, outside the active site. Such an effect may keep the ATP molecule close to the myosin surface for a sufficiently prolonged period to infer a binding event with an off-rate in the range 0.2-0.5 s^-1^. In order to directly test this idea we first blocked the myosin active site by non-fluorescent MgATP in the presence of vanadate (37) before adding Alexa-ATP (Fig. 3a-c). If the existence of the unexplained phase reflects a dynamic switch between different myosin conformers, i.e. myosin conformations with different catalytic activity upon substrate binding to the active site, it seems inevitable with a reduced amplitude of the unexplained phase after active site blocking. On the other hand, no such reduction is, a priori, expected if the unexplained phase reflects non-specific binding outside the active site. In agreement with the latter prediction, active site blocking did not reduce the amplitude of the 0.2-0.5 s^-1^ phase (Fig. 4a-c) despite substantial reduction in amplitude of the slower phase, attributed to basal MgATP turnover. In contrast, there was an increase in the absolute number of events per myosin molecule and time (legend, Fig. 4) attributable to the unexplained phase. Additionally, a new faster phase emerged. We postulate that this reflects different non-specific ATP binding properties in different myosin states. Finally, we present evidence against contribution from trivial complicating factors to the unexplained phase e.g. non-specific binding of the Alexa moiety to myosin/surfaces (Fig. S5), Alexa-ADP binding to the active site (Fig. S9) or remaining surface binding of Alexa-ATP (Fig. S5). Furthermore, complicating effects due to two myosin heads in HMM are excluded by the similarity between dwell-time distributions with S1 (with only one head) and HMM (Figs. 1, 4, S6-S7).

**Figure 4.**
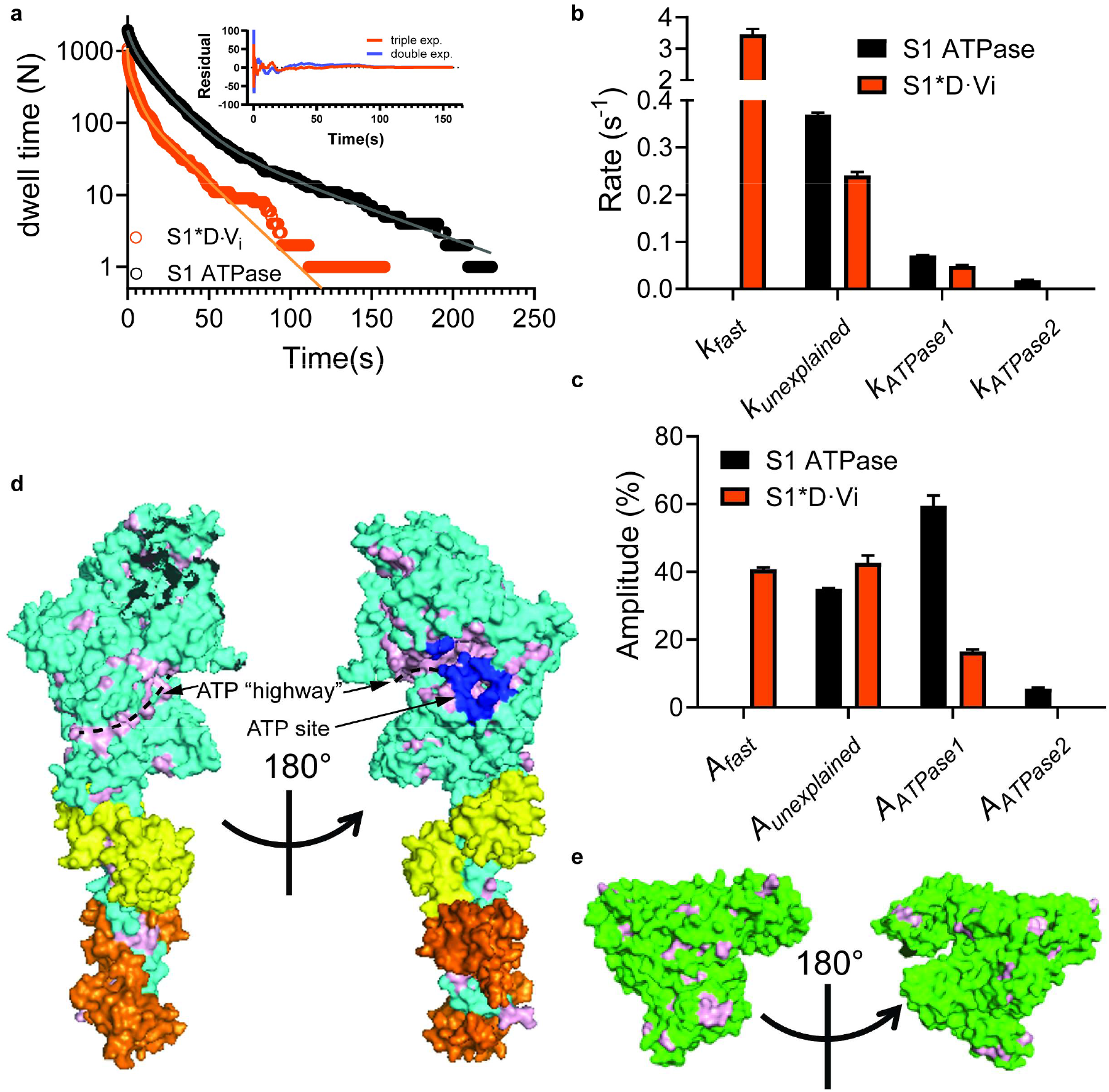
Origin of “unexplained” (0.2-0.5 s^-1^) exponential phase. **a,** Cumulative frequency distributions of Alexa-nucleotide dwell-time events on myosin subfragment 1 (S1) surface hot-spots (simple deposition of S1, Fig. 1a, sub-panel i). Data either **with** (N_dwell_ = 1964 from 127 S1 molecules in 15 min) or **without** (N_dwell_ = 1084 events from 30 molecules in 30 min) **nucleotide pocket blocking** by non-fluorescent ATP and Vanadate (S1*D·V_i_). The data were fitted by triple exponential functions (solid lines) instead of double exponential function as justified by residual plots (inset; S1*D·V_i_ data), difference in Akaike’s information criterion - AICc (4290) and R^2^ (0.9907 vs 0.9977). **b,** Rate constants obtained from fittings to data in a. **c,** Amplitudes obtained from fitting the data in a. Note that amplitude (A_ATPase1_) of basal ATPase activity (k_ATPase1_ ~ 0.05 s ^-1^) was significantly reduced for S1*D·V_i_. Unexplained phase with amplitude A_unexplained_ and rate k_unexplained_, was not appreciably affected. Note, further that fitting of S1*D·V_i_ data set produced an even faster phase (k_fast_ ~ 3.5 s^-1^). The extra slow phase with amplitude A_ATPase2_ ~ 5 % and rate k_ATPase2_ ~ 0.02 s^-1^ for the S1 ATPase may be attributed to reduced basal S1 ATPase due to S1 attachment with the head to the surface(19). Error estimates refer to 95 % confidence intervals obtained in the fits. Temperature: 23 °C. **d,** Model of S1 with predicted surface availability for ATP binding as calculated using ATPint. Total predicted surface availability for ATP binding (23.7 %, pink) vs active site (blue, 1.2 %). Proposed “ATP highway” illustrated with dashed line. **g,** Model of BSA with predicted surface availability for ATP binding as calculated using ATPint. Total predicted surface availability for ATP binding (13 %, pink). Note that unspecific ATP binding to BSA has been experimentally observed (see text).

The hypothesis of non-specific ATP binding outside the active site is supported by results using the bioinformatics tool ATPint (38) (Fig. 4d-e). Remarkably, this tool predicts substantial ATP interactions on the myosin head surface outside the active site (Fig. 4d) (11), attributing only 8 % of the ATP binding interface to the active site itself. A possible physiological role of such non-specific ATP binding is that it provides “highways” funnelling ATP to the active site by faster one-(two-)dimensional compared to three-dimensional diffusion, reminiscent of certain forms of channelling of reaction intermediates over enzyme surfaces in coupled enzymatic reactions (39). This idea accords with the non-specific binding sites surrounding, and seemingly radiating out from the active site (Fig. 4d). One may wonder whether increased rate of ATP binding could be physiologically relevant when the binding rate nevertheless is diffusion limited with a rate constant of ~10,000 s^-1^ at ~1 mM MgATP. Notably, however, actin and myosin would slide more than 1 nm relative to each other during the associated waiting time under physiological velocity of >10,000 nm s^-1^ thus imposing an appreciable non-productive braking force. Above, we hypothesized that the degree and kinetics of non-specific ATP binding may vary between myosin states. If the binding accelerates ATP transport to the active site, it would be physiologically most important in the nucleotide free actin-bound, “rigor” state. This idea fits with our observation of more frequent Alexa-ATP binding to actomyosin than to myosin. Under this interpretation, a fraction of the observed fast binding events with actin present (Fig. 1c, iii) might also represent non-specific ATP binding without active site access.

### Effect of para-aminoblebbistatin (AmBleb) on myosin basal ATPase examined on single molecule level

The optimized single molecule ATPase assay enabled us to directly observe effects of the myosin inhibitor para-aminoblebbistatin, a light insensitive, non-fluorescent and highly, soluble blebbistatin analog (Fig. 5a) (40, 41). In this proof of principle drug testing experiment, we used (Fig. 5b) 40 μM AmBleb, that produced half-maximal inhibition of the IVMA sliding velocity, to study MgATP turnover by HMM. These studies reveal appreciably longer dwell times in the presence of drug (Fig. 5c, d). For that reason, we could not use the criteria for “hotspots” described above (where we considered spots as hotspots when they pose at least 10 events per 15 min trace) and we analysed the datasets without this criterion. This shows that certain changes in experimental conditions due to drugs or the presence of mutations call for new and preferably more general criteria for hotspots. Cumulative dwell-time distributions of basal ATPase activity were best fitted using two exponential (control) and three exponential functions (AmBleb) revealing an additional slow phase of 0.005 s^-1^ in the presence of AmBleb (Fig. 5e). The other notable observation is that the rate and amplitude of the “unexplained” (“medium” in Fig. 5) process was little affected by AmBleb despite appreciable effects on the catalytic activity. Overall, these findings accord with the idea of non-specific binding of MgATP outside the active site

**Figure 5.**
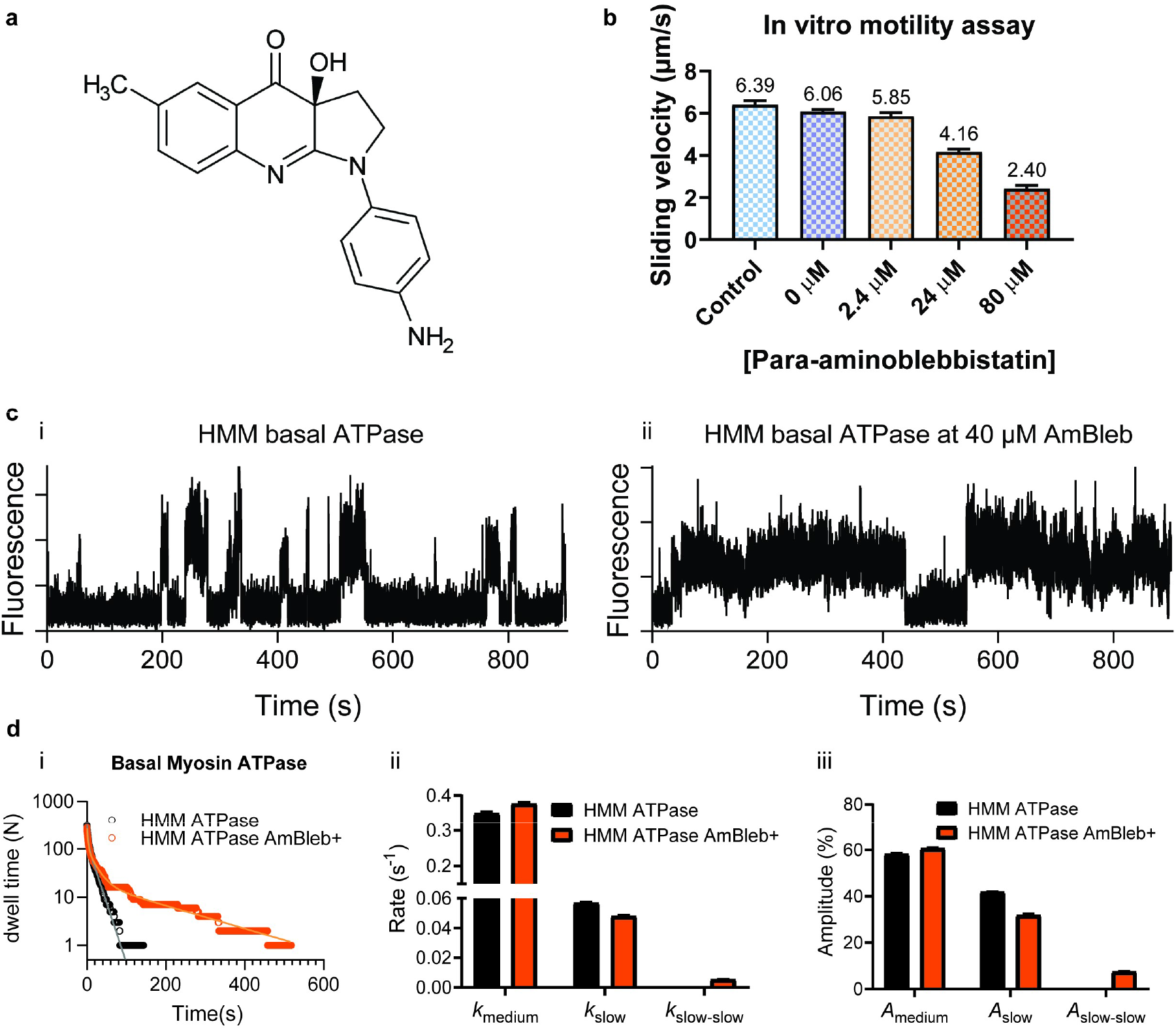
Effect of AmBleb on single molecule ATPase. **a,** The molecular structure of para-aminoblebbistatin (AmBleb) is shown. **b,** Concentration-response curve for effect of AmBleb on sliding velocity at 60 mM ionic strength (1 mM MgATP) is shown using HMM. **c,** Representative time traces of single molecule basal HMM ATPase without (i) or with 40 μM AmBleb (ii). Note two long dwell time events as direct consequence of effect of AmBleb on basal ATPase activity. **d,** Cumulative dwell time distributions of basal HMM ATPase in the absence and presence of 40 μM AmBleb were best fitted to double (control, data from 19 HMM molecules, N_dwell_ = 318) or triple (AmBleb, data from 31 HMM molecules, N_dwell_ = 280) exponential functions revealing an additional slow phase with rate constant 0.006 s^-1^ (~8 %) in the presence of AmBleb. Temperature: 20 °C.

In conclusion, we report optimizations for increased reliability and reproducibility (Figs. S2, S4, S5, S7, S8, S9) as well as expanded general usefulness of fluorescence based single molecule assays. The new insights into ATP turnover by myosin in the presence and absence of actin, using 1-100 isolated enzyme molecules (Figs. 1g-i, 3, 4), illustrate the increased capabilities. The improvements will benefit a growing spectrum of single molecule fluorescence based techniques for studies of actin and myosin, e.g. single molecule FRET (42), fluorescence polarization (43) and use of nanofabricated surfaces (e.g. zero-mode waveguides) for enhanced fluorescence background suppression (44, 45). Finally, the work paves the way for reliable single molecule analysis in high-throughput screening of drug candidates (cf. Fig. 5) with minimal requirement of proteins from costly expression systems, individual cells or patient samples.

## Materials and Methods

### Proteins

Rabbits for myosin and actin preparations were kept and sacrificed according to procedures approved by the Regional Ethical Committee for Animal experiments in Linköping, Sweden, reference number 73-14. Actin and myosin were prepared from fast skeletal muscle of New Zealand white rabbits immediately after sacrifice. Actin was prepared as described earlier (46, 47). Myosin, heavy meromyosin (HMM) and papain subfragment 1 (S1) were prepared following published protocols ((48), with modifications in (46, 49)). Protein preparations were characterized by sodium dodecyl sulfate polyacrylamide gel electrophoresis (SDS PAGE) with respect to purity as described previously(50) and concentrations were determined spectrophotometrically.

### TIRF Microscopy

We used an objective type total internal reflection fluorescence (TIRF) microscope for all single molecule experiments. The temperature was 20-25 °C but constant to within 1 °C-during a given experiment; see Figure legends. The ionic strength was 60 mM unless otherwise stated. For details, see SI Methods.

### Dwell Time Assays

To follow binding and dissociation of nucleotide, we used the Alexa647–ATP as the substrate for myosin. For detailed description of procedures for the dwell time assays, composition of assay buffer, selection criteria for signals of individual molecules and dwell time analysis please see SI Text.

## Author Contributions

A.M. and M.U. conceived the project, M.U. and A.M. designed the TIRF system. M.U. and A.M. designed the experiments. M.U. built the TÌRF system and performed single molecule fluorescence experiments. M.U. L.M. and A.M. analyzed TIRF data, L.M. performed the bioinformatics modelling. V.V. performed in vitro motility assay experiments with and without AmBleb and analyzed the data, and A.S. analyzed in vitro motility assay data, A.M., M.U., L.M., V.V. and A.S. wrote the manuscript and approved the final version.

## Acknowledgments

This work was funded by European Union Horizon2020 FET Program under contract 732482 (Bio4comp). Further, funding is acknowledged from The Swedish Research Council (grant # 2015-05290) and The Faculty of Health and Life Sciences at The Linnaeus University. Drs A. Malnasi-Csizmadia, S. Tågerud, M. Norrby and M.A. Rahman are acknowledged for valuable discussions and suggestions.

