## Supplemental information - appendix for "Optimized single molecule fluorescence sheds light on elusive enzymatic mechanisms"

by

Marko Usaj, Luisa Moretto, Venukumar Vemula, Aseem Salhotra and Alf Månsson

Supporting Information

### MATERIALS AND METHODS

#### Materials

Para-aminoblebbistatin (AmBleb) was from Optopharma. Rhodamine Phalloidin (cat. no. R415), Alexa Fluor647-Phalloidin (cat. no. A22287), Alexa Fluor647-ATP (cat. no. A22362), Alexa Fluor647-cadaverine were obtained from Thermo Fisher Scientific. Trolox (cat. no. 238813), cyclooctatetraene (COT, cat. no. 138924), 4-Nitrobenzyl alcohol (NBA, cat. no. N12821), pyranose oxidase (POX, cat. no. P4234), bovine serum albumin (BSA, standard purity, cat. no. A2153), bovine serum albumin (BSA, high purity, cat. no. A0281), dithiothreitol (DTT), glucose oxidase (GOX, cat. no. G2133), catalase (cat. no. C100), creatine phosphate (PK, cat. no. P7936), adenosine triphosphate (ATP, cat. no. A2383), creatine phosphokinase (CPK, cat.no. C3755), MOPS, KCl, $MgCl_2$, $K_2EGTA$, HCl, KOH, Liquinox, Ethanol, Methanol, Glucose, DMSO, Sodium Orthovanadate were purchased from Sigma Aldrich (now Merck). Other biochemical reagents were of analytical grade and purchased from Sigma Aldrich.

#### Buffers and solutions

Stock solutions of chemicals were prepared according to user manuals or published literature. In order to minimize solvent concentrations in the final assay solution, COT and NBA were prepared together in DMSO as 200 mM stock, aliquoted and kept at -20 °C. Trolox was prepared on the day of the experiment in a low ionic strength solution (LISS, see below) at final concentration of ~2 mM. Due to poor solubility of Trolox in water, the powder was first dissolved in methanol as a 100 mM solution and subsequently diluted with LISS buffer(1). pH was adjusted to 7.4 by KOH titration. The Trolox solution was filtered (0.2 µm) into a 10 cm Petri dish and exposed to UV-light (254 nm) to form Trolox-Quinone(1). Our optimized exposure at 120,000 µJ/cm$^2$ for 15 min using Stratalinker1800 (Stratagene) yielded ~20 % Trolox-Quinone in solution (molar ratio). Finally, a Trolox-Trolox/Trolox Quinone mixture in LISS (***TX/TQ-LISS***) was degassed for use in assay solutions (see below).

A low ionic strength solution (LISS; pH 7.4) contained 10 mM MOPS, 1 mM $MgCl_2$ and 0.1 mM $K_2EGTA$. A wash buffer was prepared to contain 50 mM KCl and 1 mM DTT in LISS. Standard assay solution(2, 3) (IVMA buffer) for gliding in vitro motility assays was prepared in LISS with addition of (final concentrations) 10 mM DTT, 45 mM KCl, 3 mg/ml glucose, 0.1 mg/ml glucose oxidase, 0.01 mg/ml catalase, 2.5 mM creatine phosphate, 0.2 mg/ml creatine phosphokinase and 1 mM MgATP. ***Optimized TIRF assay solution (TIRF buffer)*** was prepared in TX/TQ-LISS (see above) with 45 mM KCl, 10 mM DTT, 7.2 mg/ml glucose, 3 U/ml POX, 0.01 mg/ml catalase, 2.5 mM CP, 0.2 mg/ml CPK, 2 mM COT, 2 mM NBA, ~2 mM TX/TQ and 0.64% methylcellulose with or without Alexa647-ATP (up to 10 nM). Note, it is important first to add COT and NBA to TX/TQ-LISS and vortex to ensure proper solubility of these two components. Ionic strength of assay buffer was 60 mM.

The traditionally used oxygen scavenger system based on glucose oxidase - GOC (glucose/glucose oxidase/catalase) was replaced by pyranose oxidase – POC (glucose/pyranose oxidase/catalase) with the aim to prevent acidification of the assay buffer under prolonged observation(4). Catalase and pyranose oxidase were prepared in LISS as 100X solution followed by centrifugation (10,000×g, 1 min) and storage of the supernatant at 4 °C for up to four weeks.

Methylcellulose was prepared at 1.6 % (w/v) in TX/TQ-LISS, stirred overnight at 4 °C, degassed, aliquoted and stored at -20 °C.

AmBleb was prepared in DMSO, aliquoted and stored at -20 °C. The stock concentration was checked by absorbance measurement on spectrophotometer (UV1800, Shimadzu) using an extinction coefficient of 6860 $M^{-1}cm^{-1}$ (aqueous solution at pH = 7.3) according to the manufacturer´s instructions. DTT (1 M) was prepared freshly on the day of the experiments in LISS. Activated sodium orthovanadate (vanadate further in text) was prepared in water by repetitive steps of adjusting pH-boiling-cooling(5, 6). The activation was monitored by observing change of the solution color from yellow to colorless. The final pH of the 52.5 mM sodium orthovanadate stock (as determined spectrophotometrically using extinction coefficient $\varepsilon$ = 3,550 at 260 nm) was stabilized at pH 8.0, aliquoted and stored at -20 °C.

**TIRF Microscopy**

We used an objective type total internal reflection fluorescence (TIRF) microscope for all single molecule experiments. The TIRF microscope was custom-built around a Nikon epifluorescence inverted microscope Nikon Eclipse TE300 arranged on an optical table (CleanBench 63-574, TMC). A 60X oil immersion objective lens with high numerical aperture was used (CFI Apochromat TIRF 60XC Oil, NA 1.49, Oil; Nikon) and images were recorded using an EMCCD camera (Andor iXon Ultra 897 EMCCD). The frame rate was limited to 20 $s^{-1}$ to allow sufficiently long continuous recording (900-1800 s). Pixel size of acquired images was 0.267x0.267 $\mu m^2$. The beams of the red lasers (Melles Griot, 56RCS/S2799, OEM diode laser

45 mW, 642 nm or Melles Griot 05-LHP-925 30 mW, 632.8nm HeNe laser) were used to excite Alexa647.

The optical set-up is based on published design criteria(7, 8). Speckle across the field of view was minimized, using a vibrating optical fiber, to provide as uniform illumination as possible. The laser beam was coupled into the optic fiber that was rolled around the vibrator unit. The latter was realized either as an electric toothbrush (Oral-B Pro60C) or, more effectively, as a "hanging" vortex mixer (Vortex Genie 2, Scientific Industries) with custom-adapted head, vibrating at maximal speed.

Our TIRF-microscope is depicted in Fig. S1. The laser beam was coupled into the multimodal optical fiber (MMF, M14L02 - Ø50 µm, 0.22 NA, SMA-SMA Fiber Patch Cable, Low OH, 2 Meters, ThorLabs), mounted on a fiber holder (H1, SM1SMA fiber adaptor, ThorLabs), with the help of a focusing lens (L1, AC254-030-A-ML, ThorLabs). The Optical fiber was guided around the fiber vibrator (see above) and mounted on a Fiber Port (PAF-SMA-5-A, ThorLabs). With the help of a collimating lens (L2, f = 250 mm, ThorLabs) the beam was collimated and finally, using the lens L3 (AC254-250-A-ML, ThorLabs), it was focused to the objective back focal plane. Using a manual mirror, M3 on a 1D stage we achieved TIRF illumination by positioning the beam on the outer edge of the back focal plane. The same stage allows removal of the mirror from the optical path to achieve epifluorescence illumination (100 Watt Mercury Lamp) for imaging of Rhodamine phalloidin labeled actin filaments. Careful alignments of the optical elements was aided by power measurements using the USB Power and Energy Meter device (PM100USB, ThorLabs) equipped by Silicon Power Head (400-1100 nm, 50 mW, ThorLabs), controlled by dedicated software provided by the manufacturer. Maximal power measured at the back focal plane position was ~2.7 mW (diode laser) or ~0.7 mW (NeHe laser). We were able to reduce power stepwise to ~25 %, ~13% or ~4 % of maximal power by the use of Neutral Density filters. To achieve satisfactory signal to noise ratio (SNR), without noticeable increase in the bleaching rate, the experiments were performed at maximal power (NeHe laser) or at ~0.7 mW power (diode laser, ND filter in use) unless stated otherwise. In our experimental set-up, the illuminated area was disc-like with approximately 40 µm diameter. Filter cubes were used to select a suitable wavelength range according to the emission spectra of the fluorophores (Cy5 for Alexa 647 and Cy3 for Rhodamine; Omega Optics).

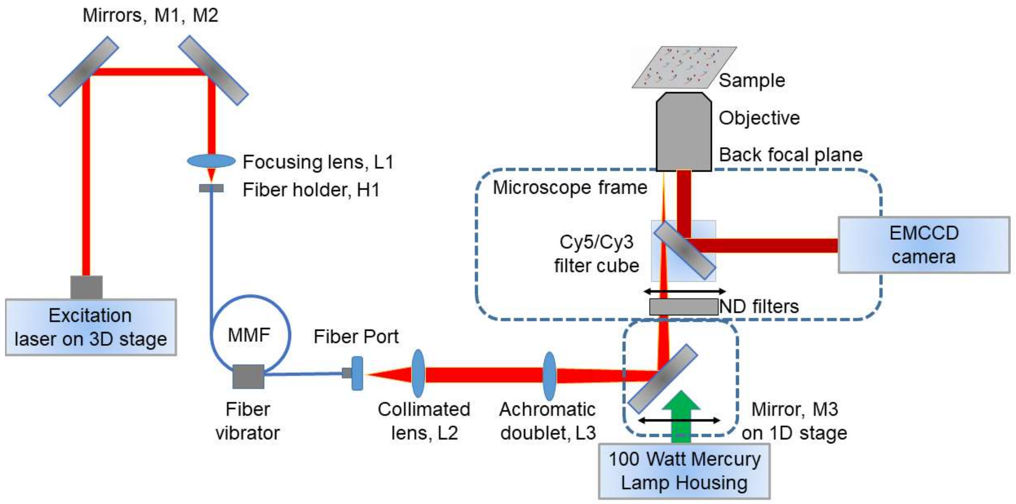


**Fig. S1.** Schematic of set-up for TIRF and EPI fluorescence microscopy.

### Surface preparations

Glass coverslips used for single molecule assays need to be particularly clean. Out of the box coverslips usually contain numerous unidentified fluorescent objects (UFOs, Fig. S2a). However, it seems that the density of UFOs and their behavior varies in a random fashion from coverslip to coverslip and from batch to batch. We could not identify the sources of the UFOs but most of commercial cover slips are flame-polished that could potentially generate nanoparticles with blinking behavior. The coverslips were cleaned in several steps. Coverslips were placed in water-filled containers, one by one and then subjected to: 20 min sonication in Liquinox (1%) and ethanol (95 %), 20 min incubation in Aqua regia and 20 min sonication in KOH (2.5 M)(9). Between different steps, coverslips were rinsed in water (3x). Aqua regia was prepared by mixing two parts of nitric acid with one part HCl(9). Subsequently, coverslips were further derivatized with trimethylchlorosilane (TMCS) as described earlier(10, 11). Notably, this includes incubation of coverslips in piranha solution (5 min, 80 °C), which per-se did not efficiently remove UFOs. *Caution! Piranha solution is a highly corrosive acidic solution, which can react violently with organic materials. Do not store in closed container, and use appropriate safety precautions. Similarly, aqua regia solutions are extremely corrosive and may result in explosion or skin burns if not handled with extreme caution. Before disposal, the solution should be cooled down and neutralized with sodium bicarbonate. Never store aqua regia in a closed container.* To passivate surfaces, high purity BSA was centrifuged

(220,000×g, 15 min) before use(12). This process yielded clean functionalized glass coverslips with quite few remaining UFOs (Fig. S2b). These were the type of slides that were used in our single molecule assays of ATP turnover. Fluorescence trajectories from remaining UFOs and non-specific Alexa-ATP binding to the surfaces are depicted in Fig S2c and Fig. S2d, respectively. The dwell times were extracted (as described further in text below) and plotted as cumulative distributions that can be fitted by double exponential functions (Fig. S2e-g). Surface preparations as described above, minimized the presence of UFOs and Alexa-ATP nonspecific binding. Notably, none of the analyzed traces from UFOs in Fig. S2e-f from properly prepared surfaces would pass the criteria for "hotspots" (see further text below) of having more than 10 events per 15 min trace.

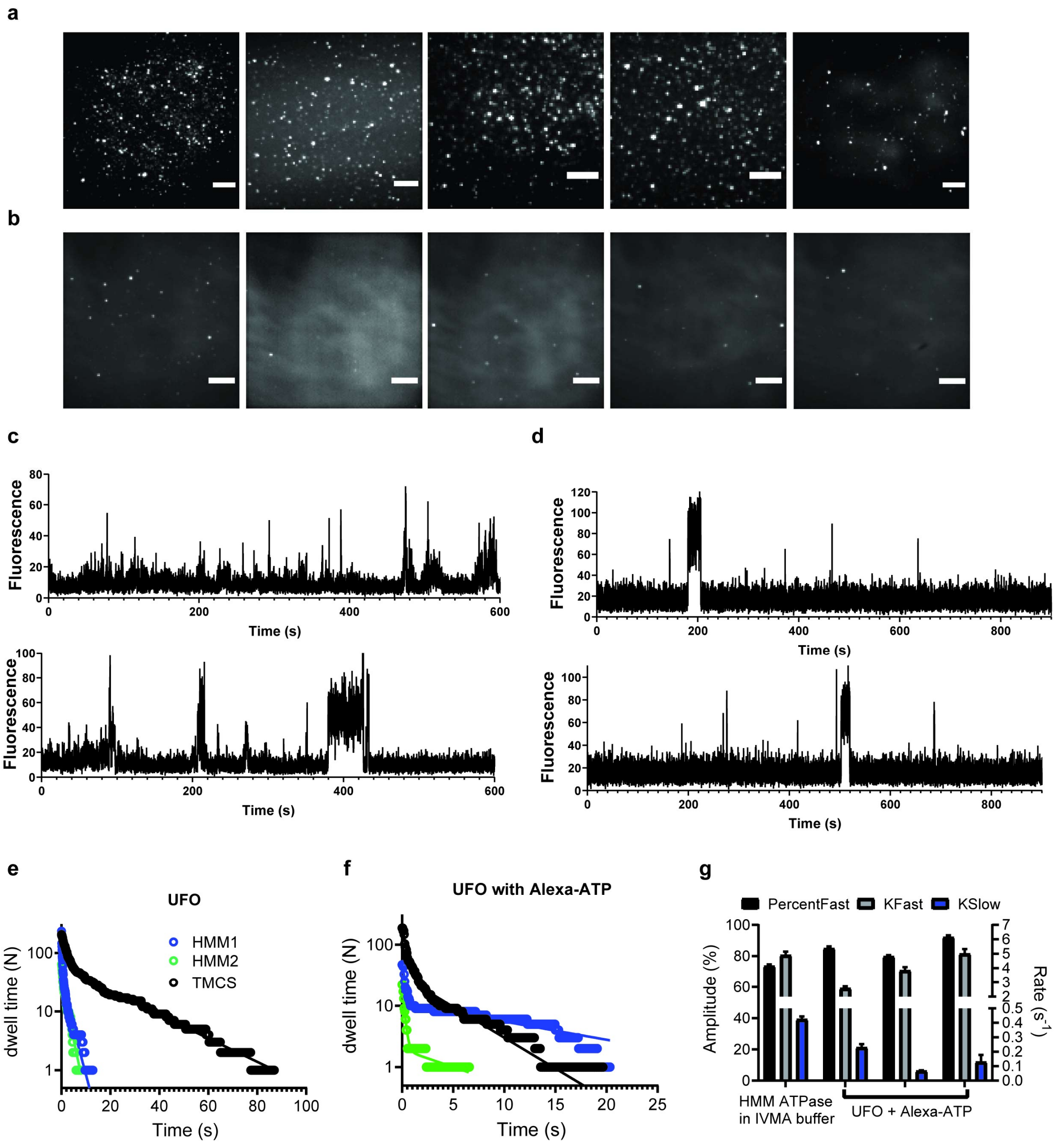

**Fig. S2. Cleaning of glass coverslip surfaces and selection of BSA.** Time projections (Fiji(13)) and analyses of 15 min videos (50 ms exposure time/frame) from different glass slides recorded in TIRF assay buffer containing 5-10 nM Alexa647-ATP. Bars, 5 μm. **a.** Out-of-the-box glass coverslips, cleaned with piranha solution, silanized and coated with standard grade BSA. **b.** Glass coverslips cleaned using optimized multistep procedure described in Materials and Methods followed by cleaning with piranha solution, silanization and coating with selected, non-fluorescent BSA. **c.** Two representative time traces of UFO fluorescence without Alexa-ATP. **d.** Two representative time traces of UFO fluorescence on non-fluorescent high-purity BSA coated surfaces with Alexa-ATP present. **e.** Cumulative dwell time distributions of UFOs (note, no Alexa-ATP present) from piranha cleaned surfaces derivatized with TMCS only ("TMCS") or also subsequently incubated with 1 nM HMM and 1mg/ml standard BSA ("HMM1" ), or with 34.3 pM HMM and 1mg/ml non-fluorescent BSA ("HMM2"). Solid lines are fits to double exponential functions. The basis for the different appearance between TMCS and HMM-labelled plots could be effects of proteins on photophysics of UFOs or just random effects. Detailed analysis is outside the scope of the study. **f.** Cumulative dwell time distributions of UFOs from non-fluorescent BSA-coated and optimally cleaned surfaces in the presence of Alexa-ATP (≤ 10 nM; three surfaces were examined) showing different behavior (seemingly random) from slide to slide. Solid lines are fittings to double exponential functions. **g.** Rates and amplitudes obtained from fitting the data in f. Note how the values can be in the similar range as seen for HMM ATPase studied in an IVMA buffer (Fig. 1f, main paper). However, importantly, the bright spots to which data in Fig. S2f-g refer, would not pass the criteria for "hotspots" (at least 10 binding events per 15 min trace; see further text) to be included in studies of ATP turnover.

### In vitro motility assays

The experiments were performed using flow-cells assembled with trimethylchlorosilane derivatized glass coverslip (#0 or #1, 24 × 60 mm) for the floor and untreated glass coverslip (#0, 18 x 18 $mm^2$) for the ceiling of the cell, spaced (~100 μm) with double-sided tape (3M Scotch). The flow cell volume was 10-15 μl.

For in vitro motility assays(3, 14), flow cells were incubated with HMM (120 μg/ml; 5 min) and then with BSA (1mg/ml; 2 min), both in Wash buffer. Subsequently, the flow cells were rinsed with Wash buffer, incubated with 2-10 nM rhodamine-phalloidin labeled actin filaments (prepared in Wash buffer) and rinsed again with Wash buffer, prior to addition of the assay solution (IVMA buffer). Image acquisition was performed using an inverted fluorescence microscope Zeiss (Axio Observer.D1, Zeiss, Germany, with a 63x objective, NA=1.4). For excitation, a Mercury short-arc lamp (OSRAM GmbH) was used together with suitable filter sets allowing observation of Rhodamine fluorescence. Image sequences were recorded using an electron multiplying charge-coupled device (EMCCD) camera (C9100-12PHX1, Hamamatsu Photonics) with pixel size (on sample) of 0.16x0.16 $\mu m^2$ and 0.24x0.24 $\mu m^2$ for 100x and 63x objective, respectively. The actin filament movements were recorded using a frame rate in the range 4-10 frames/second. Experiments were performed within the 23-25 °C temperature range but temperature was constant to within 1 °C during a given experiment (see further figure legends for details). Actin filament gliding velocities were calculated as described earlier(2, 15). The cut-off of the coefficient of variation (CV) (standard deviation of frame-to-frame velocity divided by average velocity in ten frames) for data inclusion in velocity analysis was set to 0.2.

**Alexa647-ATP**

We selected Alexa647–ATP as fluorescent ATP analogue because of extensive previous characterization in the lab(16) showing similar sliding velocities in the IVMA and similar basal HMM-catalyzed MgATP turnover as standard non-fluorescent MgATP(16, 17). Furthermore, importantly, all four isomers of Alexa647-ATP were found to behave similarly with respect to basal myosin ATPase. Finally, the creatine-kinase/creatine phosphate ATP-regenerating system seems to be compatible with Alexa647-ADP(18) which is useful in preventing accumulation of the latter product. This is in contrast to Cy3-labeled nucleotides which are not suitable substrates for the ATP-regenerating system(19). On the other hand, and in contrast to Cy3–ATP, the fluorescence intensity of Alexa647-ATP is reduced upon binding to myosin(16).

**Photobleaching of Alexa647**

The single molecule photobleaching rate of Alexa647 was determined using two assays: either actin filaments sparsely labeled with Alexa647-phalloidin (molar ratio Alexa647- phalloidin : actin filaments = 1:240) or Alexa647-ATP locked into the ATP-pocket by vanadate as described previously(17). The latter approach is important as a supplement because it examines the dye photophysics in an environment similar to ATPase studies. Standard deposition of myosin (heavy meromyosin or subfragment 1)-Alexa-ATP-vanadate complexes was used for these experiments. The time traces were extracted as described above with the help of time projection images. The total bleaching time was determined from events, which produced single step decrease in fluorescence intensity. We have also noted recurrence of a high fluorescence state suggesting dye transition back to the bright state from dark state. In such cases, we collected the times of the first step decrease in fluorescence intensity only. Fitting cumulative distributions of those events yielded single molecule photobleaching rate constant. Under our optimized conditions, the latter was usually an order of magnitude slower than that for the basal myosin ATPase (Fig. 1).

**Myosin deposition for single molecule assays**

Many functional assays benefit from the use of unmodified myosin molecules, i.e. without conjugation to a fluorescent dye or expression with a fluorescent protein. In this study we therefore used non-labeled myosin constructs. For this reason, it was essential to have high confidence that collected signals originate from myosin-associated events. We deployed several strategies to increase this confidence.

a) Simple myosin deposition assay

In the simple myosin deposition, we infused myosin motor fragments in pM concentrations to allow them to adsorb in random locations on the silanized surface. This procedure (Fig. 1a-b, i) is similar to that used in all (to the best of our knowledge) studies of single molecule myosin basal ATPase until now. The myosin motor fragments at such low concentration were prepared in wash solution further supplemented with 0.1 mg/ml of BSA to prevent myosin degradation and protein loss through attachment to tube wall. Recorded videos were first subjected to simple background subtraction using Fiji (Fiji Is Just ImageJ) function "Process/Subtract Background" with parameter "Rolling ball radius" set to five, successfully decreasing noise of free dye as well diminishing any illumination non-uniformities while preserving signal features. Time projection images of background-subtracted videos were created using the Fiji function "Image/Stack/Z-project/STD". These images were then used to manually select hotspots, i.e. regions of interest of 3 × 3 pixels with significant time (defined below) of high fluorescence intensity (ON-time). Using this approach, it is essential to start with clean surfaces. We used chambers with different densities of deposited myosin molecules. As shown in Fig. 1bi, the number of fluorescent spots in a field of view approximately scaled with the concentration of myosin in the solution infused into the assay chamber for the deposition of myosin molecules. To strengthen the probability that the selected spot is truly (active) myosin we only consider "hotspots" i.e. spots with at least 10 repetitive binding events (ON-times) during a 15 min recording; a criterion used previously(20). We also employ previously suggested procedures to optimize the selection of specific binding events of individual Alexa647 molecules(20). For example, we included only those signals for which both sudden one-step increase and sudden one-step decrease occurred while signals arising from more than one fluorophore beginning and ending in double- or even multiple-steps were excluded.

b) Optimized myosin deposition assay

As a novelty, we developed and used optimized myosin deposition via actin filaments (Fig. 1a-b, ii-iii). In this approach myosin motor fragments in pM concentrations (typically 85.75 pM) were preincubated with 40 nM actin filaments (G-actin concentration) labeled with Rhodamine-phalloidin in wash buffer supplemented with 0.1 mg/ml BSA for at least 10 min on ice to form actomyosin. Actomyosin was then infused into a flow cell to allow attachment to the silanized surface (5 min, RT). Subsequently, the flow cell was incubated with BSA (1 mg/ml, 2 min) and we obtained images of the actin filaments. To measure myosin basal ATPase activity, actin filaments were washed away by incubation with non-fluorescent MgATP (~ 100 μM) in wash buffer for 2 min following extensive washing with ATP-free wash buffer. Images of any remaining actin filaments were acquired. With this procedure, actin filaments were either completely removed or small actin filament fragments remained presumably due to certain number of ATP insensitive heads (Fig. 1a-b, ii). These remaining filament fragments are potentially useful for alignments between different steps, e.g. to correct for possible drift due to media exchange, etc. Finally, assay solution (IVMA or TIRF buffer) was injected into the chamber and a video was recorded. Time projected images of background-subtracted video recordings of nucleotide binding were merged with both images of the actin filaments using a Fiji function ("Image/Color/Merge channels"). Only hotspots co-localized with actin filaments before ATP wash and without actin filaments after ATP wash were considered for analysis of myosin basal ATPase. The optimized deposition of actin filaments also gives us excellent opportunity to reliably measure single molecule actin activated myosin ATPase for the first time. To this end, the actin filaments were not washed away after deposition. Rather, a gentle wash was performed with low [MgATP] of 100 nM to remove myosin motor fragments that were attached to the actin filament but not to the surface (Fig. 1a,ii; 1b,iii). The rinsing step enhanced single molecule detection by avoiding overlapping myosin heads in single ROIs of 3 × 3 pixels. Furthermore, the procedure also blocked the active sites of any non-functional myosin heads that showed irreversible ATP binding (without turnover activity). The latter could otherwise produce unrealistically long fluorescence ON events, limited only by fluorophore bleaching. Again, images of actin filaments were acquired before and after ATP wash. Only hotspots of Alexa647-ATP binding that co-localized with both actin images were considered for analysis of actin activated myosin ATPase.

**Summary of optimized approach for TIRF based ATPase assays.**

In order to remove the effect of UFOs, flow cell surfaces were extensively cleaned as described above and optimally selected and treated BSA was used for surface blocking. Generally, and unless otherwise stated, myosin motor fragments that were used for ATPase assays were deposited together with actin filaments to give highest confidence in defining the locations of the motors. Independent of mode of deposition, the analysis was limited to surface hotspots as defined above. Further, the assay was performed using ***optimized assay solution (TIRF buffer)*** containing: TX/TQ-LISS with 45 mM KCl, 10 mM DTT, 7.2 mg/ml glucose, 3 U/ml POX, 0.01 mg/ml catalase, 2.5 mM CP, 0.2 mg/ml CPK, 2 mM COT, 2 mM NBA, ~2 mM TX/TQ (TQ/TX = 0.2) and 0.64% methylcellulose with or without Alexa647-ATP (up to 10 nM). The basis for the optimized assay procedure is described in detail and further justified elsewhere in the paper.

**Details of data Analysis**

We followed recently laid out principles for analysis of dwell-time trajectories (20-22). Software routines were written in MATLAB for semi-automated analysis of Alexa-ATP dwell time recordings and to extract time traces of each individual fluorescence spot. The first step in the software routine is to allow the user to select fluorescence spots as well as nearby background ROIs on a time projection image, created as described above. An individual ROI is defined as a 3×3 pixel area. The integrated intensity of each ROI is calculated frame by frame for the entire sequence and stored as a time series. Beside batch generation of time traces from multiple ROIs we have also used manual (ROI by ROI) extraction using the Fiji function (Image/Stacks/Plot Z-axis Profile). These time series are finally analyzed manually to measure the dwell times of fluorescence spots, i.e., the time from a one-step increase of intensity above a pre-defined threshold until the single-step drop in intensity back below the intensity threshold. Only fluorescence events that start and end with a one step change in intensity were considered. In the optimized deposition of myosin, only those regions of interest were included that co-localized with an actin filament visualized by Rhodamine-phalloidin labeling (see above). The resulting dwell times were interpreted as the time spent by Alexa647-nucleotide (Alexa647-ATP and Alexa647-ADP) bound to an immobilized myosin molecule. Collected dwell times were plotted as cumulative distributions as described before(20). The data were fitted using non-linear regression in GraphPad Prism. Reported rates and amplitudes represent mean ± 95% CI.

**Modeling of ATP binding**

The predicted ATP binding residues, obtained from the ATPint(23) tool, were plotted on the surface of rabbit fast skeletal myosin 2 (PDB: 5H53) or BSA (PDB: 3V03) using PyMOL(24). Cyan: myosin, light chains: orange and yellow, active site: dark blue, binding residues: pink,

BSA: green. Beside ATPint we have also tested other ATP binding-prediction software (e.g. ATPbind(25), TargetATPsite(26), IBIS(27)). Most tools find the active site ATP binding as well as some outside residues (these differed per tool). ATPint, however, finds all the outside residues found by other tools. Supported by the fact that ATPint could predict nonspecific ATP binding to BSA(28, 29) we selected this software for further use.

**Supplemental results and discussion**

*Optimizing the assay buffer for TIRF experiments*

A unique cocktail of triple state quenchers (Trolox, COT, NBA) have been used before to stabilize the Alexa647 fluorophore in a context-dependent manner i.e. to provide photostabilization in different dye microeviroments(30). To the best of our knowledge, however, none of these triple state quenchers have previously been used for single molecule assays of myosin or actomyosin(17). Since it is known that each fluorophore may require its own triple state quencher(s) and that the microenvironment can influence fluorophore photophysics(31) we first investigated how the different triple state quenchers affect Alexa647 in the context of myosin and actin. As can be seen from Fig. S3c, the combination of all triple state quenchers was most effective. As a novelty of this study, we also optimized the ratio between Trolox-Trolox and Trolox-Quinone with respect to the effectiveness in combating photophysical complications (Fig. S3e-f). In the process of the optimization, we exposed the Trolox solution to UV light for different times(1, 32), measured the Trolox-Trolox and Trolox-Quinone fractions spectrophotometrically (Fig. S3e)(1) and then tested the mixture in the single molecule assay. This analysis (Fig. S3e-g) suggested an optimal Trolox-Trolox/Trolox-Quinone ratio of ~0.2 that was subsequently used.

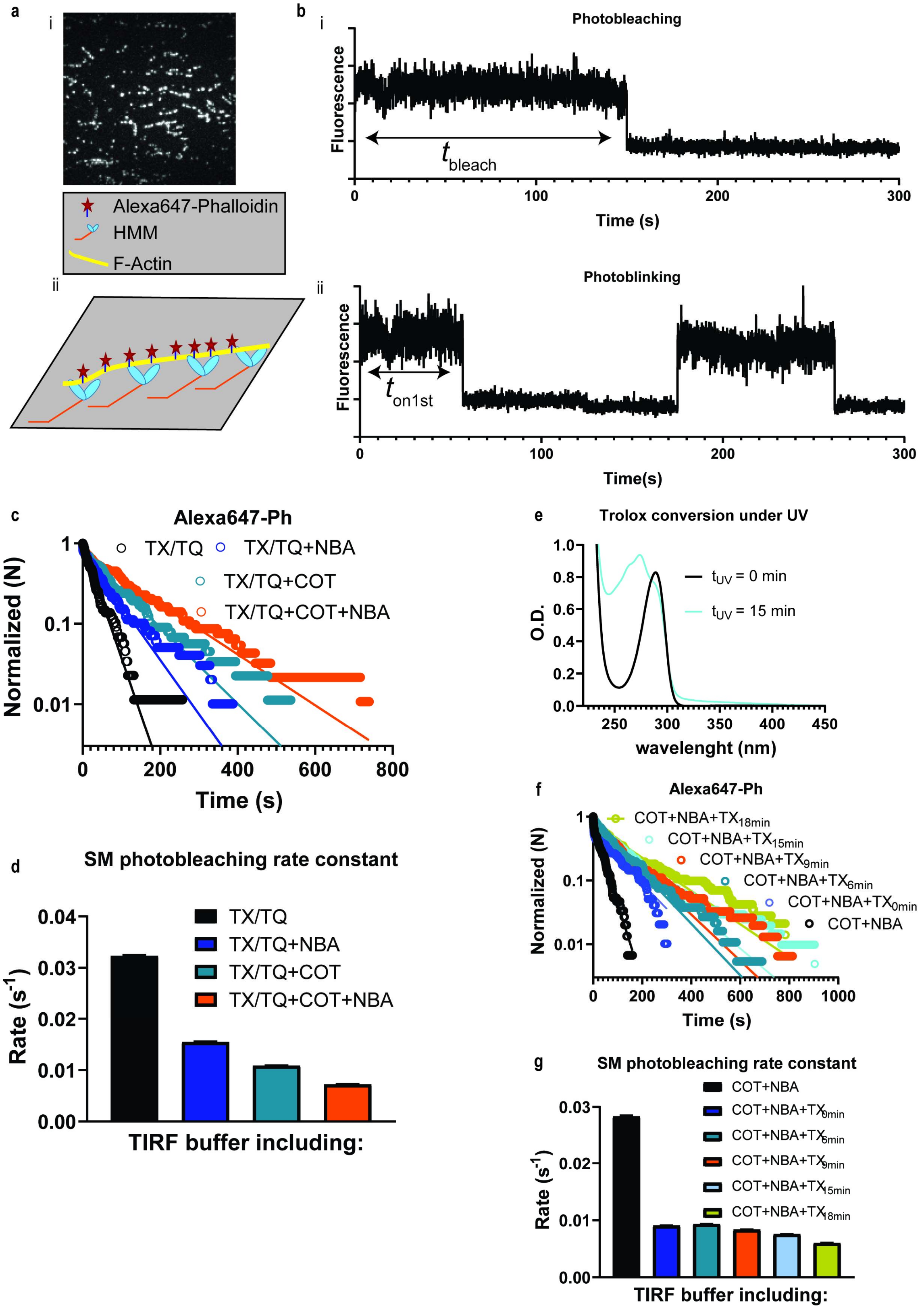
a
i
Alexa647-Phalloidin
HMM
F-Actin
ii
b
i
Photobleaching
Fluorescence
t bleach
Time (s)
ii
Photoblinking
Fluorescence
t on1st
Time(s)
c
Alexa647-Ph
TX/TQ
TX/TQ+NBA
TX/TQ+COT
TX/TQ+COT+NBA
Normalized (N)
Time (s)
d
SM photobleaching rate constant
TX/TQ
TX/TQ+NBA
TX/TQ+COT
TX/TQ+COT+NBA
Rate (s-1)
TIRF buffer including:
e
Trolox conversion under UV
tUV = 0 min
tUV = 15 min
O.D.
wavelenght (nm)
f
Alexa647-Ph
COT+NBA+TX18min
COT+NBA+TX15min
COT+NBA+TX9min
COT+NBA+TX6min
COT+NBA+TX0min
COT+NBA
Normalized (N)
Time (s)
g
SM photobleaching rate constant
COT+NBA
COT+NBA+TX0min
COT+NBA+TX6min
COT+NBA+TX9min
COT+NBA+TX15min
COT+NBA+TX18min
Rate (s-1)
TIRF buffer including:

**Fig. S3. Stabilizing Alexa647 dye photophysics. a,** Image (i) and schematic presentation (ii) of F-actin filaments partly labelled (1:240 molar ratio) with Alexa647-Ph, which was used in these experiments to uncouple fluorescence change due to ATPase activity and photobleaching/blinking. Bar, 5 µm. **b,** Representative time traces of single molecule photobleaching (i) and photoblinking (ii) of Alexa647-Ph. Photobleaching times ($t_{bleach}$) or times until first photoblinking event ($t_{on1st}$) were measured and used in further analysis below. **c,** Cumulative frequency distribution of single molecule Alexa647 $t_{bleach}$ and $t_{on1st}$ in TIRF buffer containing different triple state quenchers and redox active components: Trolox-Trolox/Trolox-Quinone (TX/TQ), 4-Nitrobenzyl alcohol (NBA), Cyclooctatetraene (COT). Note appreciable increase in $t_{bleach}$ or $t_{on1st}$ when all three components were included. The distributions are satisfactorily fitted by a single exponential function (solid lines). **d.** Single molecule photo-bleaching (-blinking) rate constants obtained from the fitting of data in **c**. Note appreciable reduction of rate constant values by performing experiments in TIRF buffer containing all three components. **e.** TX conversion to TX/TQ under exposure to UV light as observed by difference in absorbance spectra(1). **f.** Cumulative frequency distribution of single molecule Alexa647 $t_{bleach}$ and $t_{on1st}$ in TIRF buffer containing NBA and COT which was further supplemented by TX/TQ with varying amount of TQ achieved by exposing the TX solution to different amounts of UV radiation (in min). Note appreciable increase in $t_{bleach}$ or $t_{on1st}$ when [TQ]/[TX] reaches ~ 0.2 at 15 min UV exposure. The distributions are well fitted with a single exponential function (solid lines). **g.** Single molecule photo-bleaching (-blinking) rate constants obtained from the fitting of data in **c**. Note appreciable reduction of rate constant values by performing experiments in TIRF buffer containing all three components at higher [TQ]. We used 15 min UV exposure ([TQ]/[TX] ~ 0.2) for all experiments unless otherwise stated.

We also examined the possibility that different fluorophore microenvironments of Alexa-ATP in the myosin active site and Alexa-Phalloidin on actin may affect photobleaching/blinking. The studies of Alexa-nucleotide in the active site were performed by first forming a stable myosin·Alexa-ADP·Vi complex(33, 34) to allow studies of blinking/bleaching without interference from turnover events. Although the bleaching rate constants were comparable between Alexa-Phalloidin and Alexa-ATP/ADP locked in the myosin active site pocket, extra viscosity was needed (realized by adding methylcellulose) to stabilize the fluorescence signal (Fig. S4).

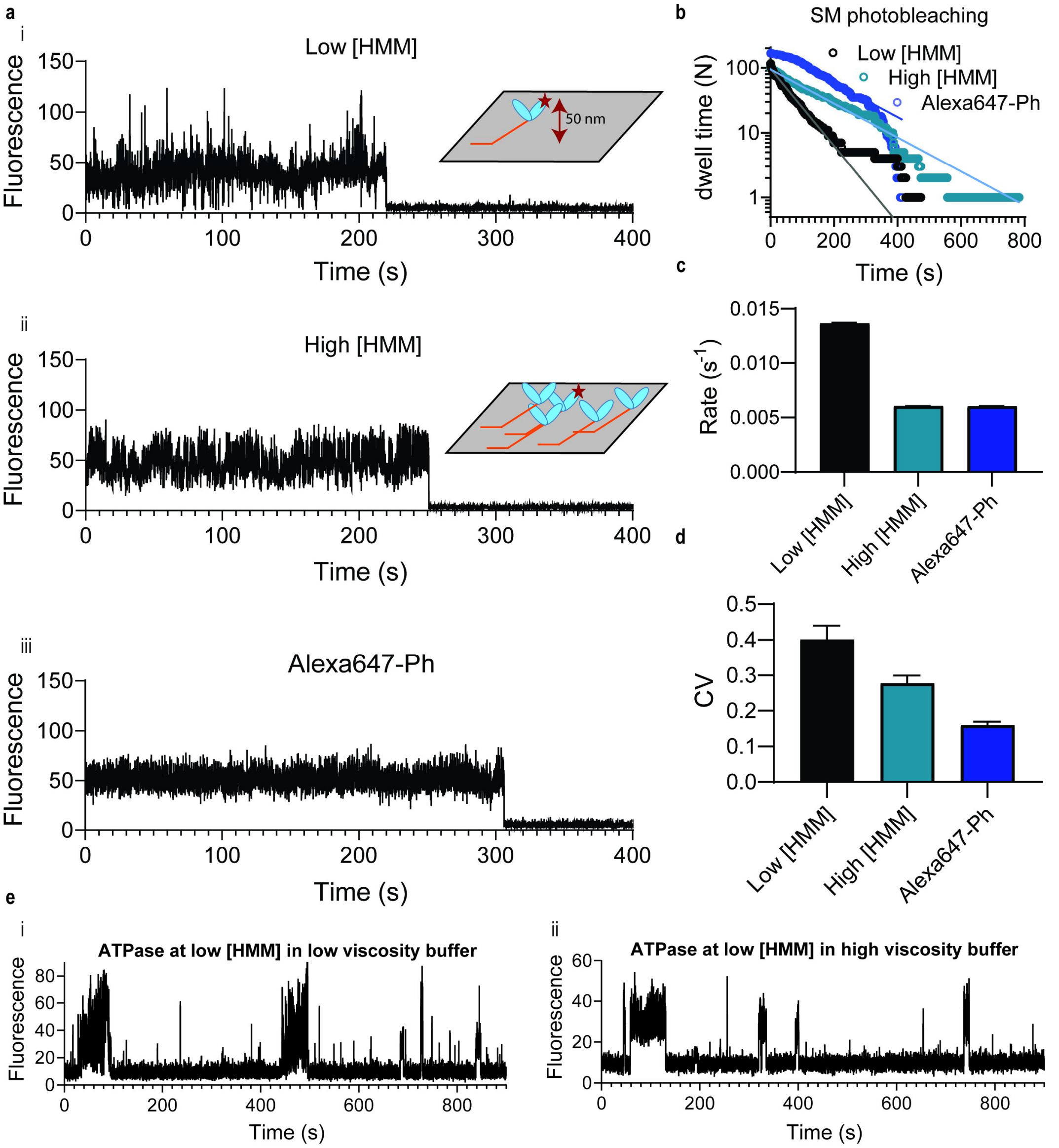


**Fig. S4. Stabilizing the Alexa647 dye signal. a,** Representative time traces for single molecule photobleaching of Alexa647-ATP locked with Vanadate into nucleotide pocket (so called HMM*D·Vi complex) at (i) low [HMM*D·Vi], (ii) at low [HMM*D·Vi] supported with saturated concentration of nucleotide-free HMM, (iii) Alexa647-Ph (as in Fig. S3). **b,** Cumulative frequency distribution of single molecule Alexa647 photobleaching under experimental conditions presented in a. The distributions can be fitted with a single exponential function (solid lines). **c,** Single molecule photo-bleaching rate constants obtained from the fitting of data in b. **d,** Signal stability estimated by calculation of signal CV. Note appreciable reduction of CV when HMM*D·Vi was stabilized by other HMM molecules or when observing the fluorophore firmly attached via phalloidin to the actin filament. Appreciable noisiness of the signal in the case of low [HMM*D·Vi] may be attributed to thermal fluctuations of HMM. Such fluctuations (~ 50 nm up and down in total, see(17)) would be sufficient under evanescent wave TIRF illumination to cause considerable fluctuations of the fluorescence signal. **e,** Representative time traces of single molecule HMM ATPase performed in TIRF buffer with (i) or without (ii) methylcellulose. High viscosity of the buffer damped thermal fluctuation of HMM, improving overall signal stability.

*Specific versus nonspecific binding of Alexa647–ATP and Alexa647 moiety to assay surface.*

By use of extensive surface cleaning, optimized selection of BSA and deposition of myosin via actin (main Fig. 1a ii-iii) we have greatly minimized the risk of artefacts due to dwell time events not associated with myosin motor fragments. However, with the aim to fully eliminate this risk we compared the intensity–time traces of regions of interest (3 × 3 pixels) *outside “hotspots*” - that is, outside regions of 3 × 3 pixels with many repetitive binding events - versus intensity–time traces of “*hotspots” themselves*. From comparison of the traces and the collected dwell times we estimated that, on average 2 ± 1 (mean ±SD, N=168 traces from 5 independent experiments) dwell times appear in a region of interest of 3 × 3 pixels throughout a total observation period (15 min) in the absence of myosin. The plotted cumulative dwell time distributions for out-of-hotspot events were differing greatly from the distributions for the hotspots (Fig. S5). For control purposes (Fig. S5c), we subtracted out-of-hotspot events from hotspot events. This was done by arranging the events from background and hotspots into two histograms with bins equal to exposure time (52 ms) followed by bin wise subtraction ($Hist_{hotspots}$-$Hist_{background}$). Fitting of the corrected cumulative dwell time distribution from hotspots yielded rates and amplitudes essentially equal to those obtained from uncorrected dwell time distributions (Fig. S5a-c). This confirms that nonspecific Alexa647-ATP binding to the surface (BSA or the underlying surface), under our experimental conditions is negligible. Furthermore, in order to examine any unspecific binding of the Alexa647 moiety of Alexa-ATP, we utilized Alexa647-cadaverine. From the results of binding experiments, designed as for the true ATPase studies except for replacing Alexa-ATP by Alexa647-cadaverine, we conclude that the Alexa647 moiety does not play any role in Alexa-ATP binding events (Fig. S5d-e).

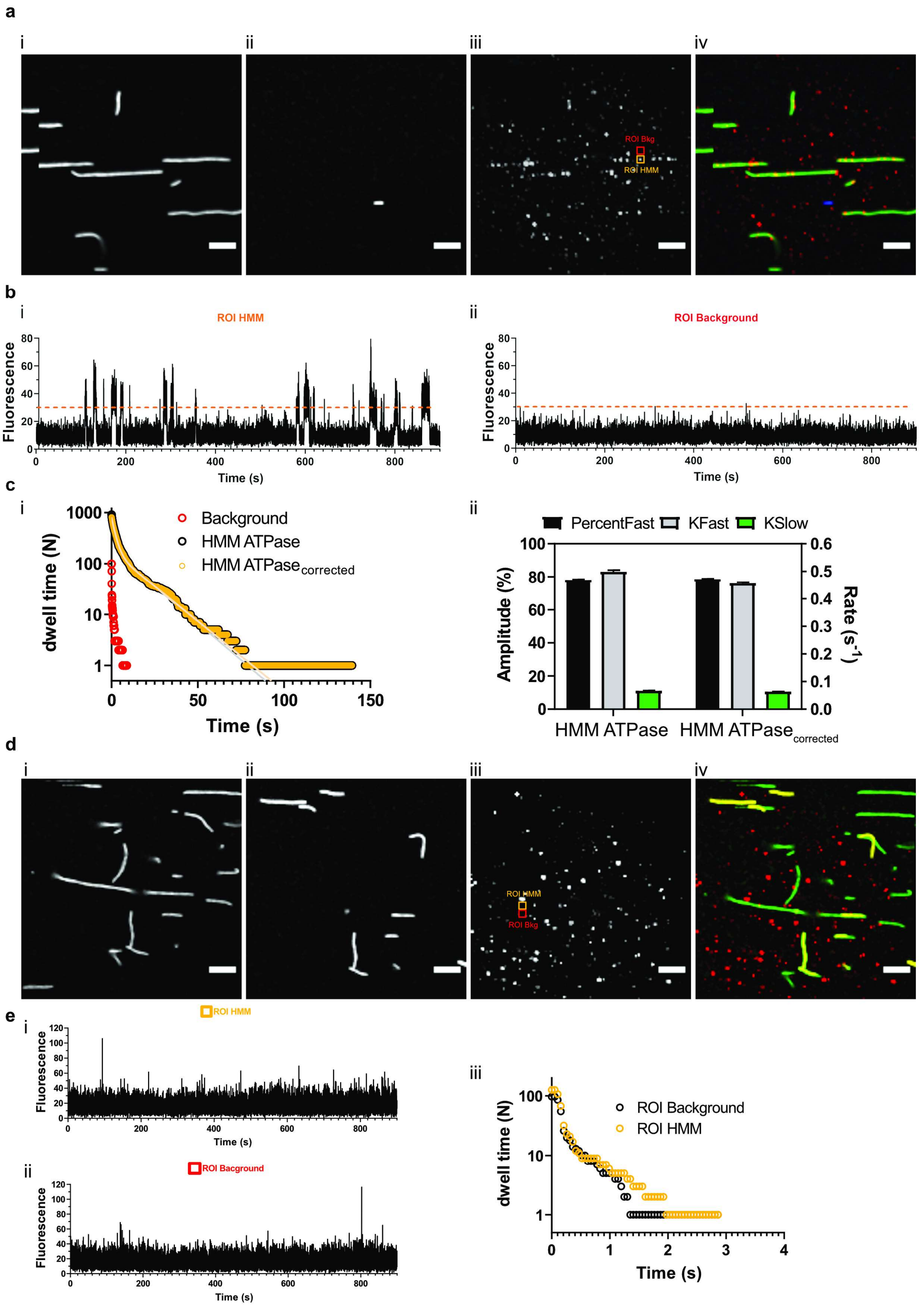

**Fig S5. Specific vs unspecific binding of Alexa-ATP and Alexa647 moiety (using Alexa647-cadaverine). a,** Figure set depicting optimized HMM deposition with Rhodamine phalloidin labeled actin filaments before (i) and after (ii) high [ATP] = 100 μM wash together with (iii)

Alexa-ATP time projection fluorescence (see text for definition) of 15 min videos (50 ms exposure time/frame). Only spots colocalized with washed-away F-actin filaments were included in the analysis (vi, green - F-actin filaments before ATP wash, blue - F-actin filament after ATP wash, red - time projected Alexa-ATP binding events). Bars, 5 µm. **b**, Representative time traces of “ROI HMM” (see a, iii) depicting HMM basal ATPase and “ROI Background” (“ROI-Bkg”, see a, iii) depicting any unspecific Alexa-ATP binding in immediate vicinity of the HMM location. The dashed line presents the same threshold used in both traces to collect dwell time events in a consistent manner. **c,** Cumulative frequency distribution of Alexa-nucleotide dwell-time events in background, HMM surface hot-spots, and HMM background-corrected hotspot (see also text). The latter two were well fitted with double exponential functions (solid lines). HMM ATPase data from 33 HMM molecules, $N_{dwell}$ = 911, were corrected using 33 background traces with $N_{dwell\text{-}background}$ = 99. After subtraction, the HMM $ATPase_{corrected}$ had $N_{dwell}$ = 813. Note, no apparent difference in rates and amplitudes after HMM hotspots were background corrected. **d**, Figure set examining nonspecific binding of Alexa647-cadaverine not expected to bind to the myosin active site. Optimized HMM deposition with Rhodamine phalloidin labeled actin filaments before (i) and after (ii) high [ATP] = 100 µM wash together with (iii) Alexa647-cadverine time projection fluorescence of 15 min videos (50 ms exposure time/frame). Only ROIs colocalized with washed-away F-actin filaments were included in the analysis (iii, green - F-actin filaments before ATP wash, yellow - F-actin filament after ATP wash, red - time projection of Alexa647-cadaverine binding events). Bars, 5 µm. **e,** Representative time traces of (i) “ROI HMM” (from d, iii) depicting events on what seems to be HMM hotspots after adding Alexa647-cadaverine and (ii) “ROI Bkg” (see also d, iii) depicting events in immediate vicinity of the presumed HMM hotspot. Note (iii) no apparent difference in dwell time distributions for events associated with actin filament (“ROI-HMM”) and not (“ROI-Bkg”). In total 58 ROIs were analyzed: 29 ROI HMM, 29 ROI Background producing 126 and 99 dwell times, respectively, similar to non-specific binding of Alexa-ATP outside hotspots (see **c** above).

*S1 and HMM ATPase under optimized conditions*

We used the optimized assay conditions (cf. summary section under Materials and Methods) to study single molecule S1 (simple deposition method) and HMM (optimized myosin deposition) basal ATPase (Fig S6). In all cases, there was a slow phase consistent with basal MgATP turnover by myosin. An almost 10-fold faster unexplained phase was also invariably present. The results were quantitatively similar in two experiments using S1 and two experiments using HMM. This demonstrates the reproducibility of the assay as well as the negligible complications introduced by the two-headed nature of HMM under our optimized assay conditions.

Since the fastest phase (rate constant >2 $s^{-1}$), ubiquitous and dominant in a previous study(20) was completely lost we tried to fit the data from Fig. S6 with triple exponential functions in order to potentially resurrect this phase. As can be seen from Fig. S7 triple exponential fits did not achieve this, despite setting initial values in the regression procedure such that the rate constants were similar to the fastest ones seen previously. Rather an extra slow phase emerged but with quite small amplitude.

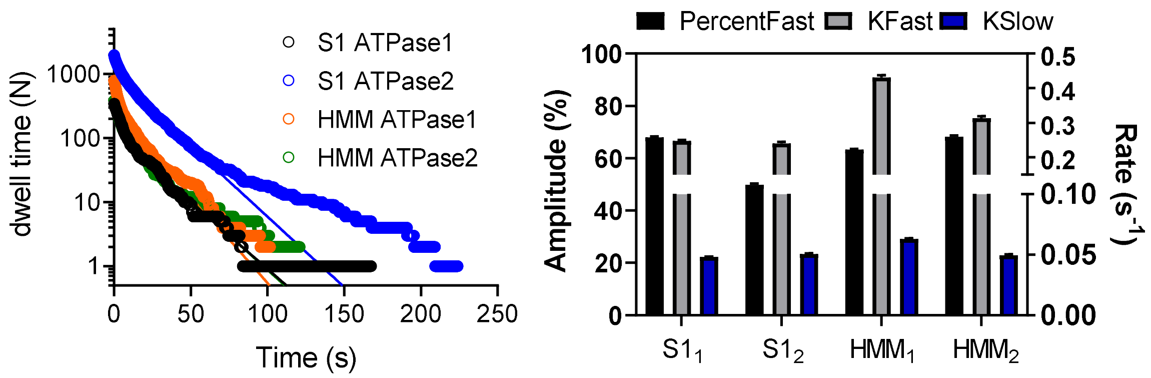


**Fig. S6. Reproducibility of optimized single molecule ATPase assay.** *Left*: Cumulative frequency distribution of Alexa-nucleotide dwell-time events comparing HMM and S1 basal ATPase activity under optimized conditions on four different experimental occasions. The distributions are well fitted with double exponential functions (solid lines). Right: Amplitudes and rate constants obtained from the fitting of data on the left. Columns represent means and error estimates refer to 95 % confidence intervals. Temperature: 23 °C.

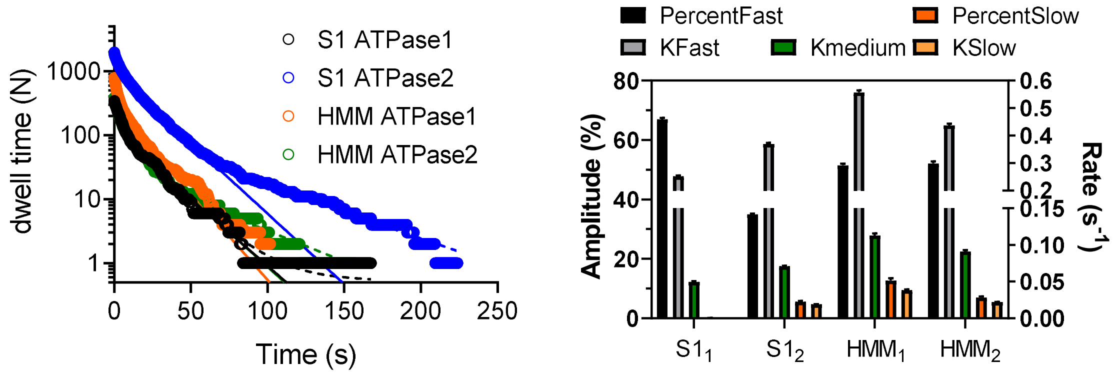


**Fig. S7. Double-exponential vs Triple exponential fit to cumulative frequency distributions for Alexa-nucleotide on-time events.** *Left*: Cumulative frequency distribution of Alexa-nucleotide dwell-time events comparing HMM and S1 basal ATPase activity under optimized conditions. The distributions are fitted with double (solid lines) or triple (dashed lines) exponential functions. *Right:* Amplitudes and rate constants obtained from the fitting of data on the left using triple exponential function. Bar heights represent mean values and error estimates refer to 95 % confidence intervals. Note, the "fast phase" corresponds to the unexplained phase in double exponential fittings whereas the phase attributed to ATP turnover in double-exponential fits is here subdivided into a "medium" (0.05-0.1 $s^{-1}$) and "slow" (<0.05 $s^{-1}$) phase. Temperature: 23 °C. Qualities of the fits and further details in Table S1.

**Table S1: Summary of best mean parameter values in double, triple, or triple with weighting (1/time) exponential fits. For 95 % CI estimation please see corresponding figures (Fig. S7, Fig S8):**

| ATPase | $k_{fast}$ ($s^{-1}$)<br>$A_{fast}$ (%) | $k_{inter}$ ($s^{-1}$)<br>$A_{inter}$ (%) | $k_{slow}$ ($s^{-1}$)<br>$A_{slow}$ (%) | $r^2$ | AICc |
|---|---|---|---|---|---|
| $HMM_1$, N=785<br>*triple exp.* | 0.56<br>51.5 | 0.11<br>35.8 | 0.038<br>12.7 | 0.9995 | 3782 |
| [a] *triple exp*<br>*weight 1/t.* | 0.3773<br>~ 10.91 | ~ 0.0586<br>~78.18 | ~ 0.0583<br>~ 10.91 | 0.9992 | 5233 |
| *double exp.* | 0.43<br>63.29 | 0.063<br>36.71 | | 0.9985 | 6064 |
| $HMM_2$, N=387<br>*triple exp.* | 0.44<br>52.2 | 0.09<br>40.8 | 0.022<br>7.0 | 0.9991 | 2580 |
| [a] *triple exp*<br>*weight 1/t.* | 0.275<br>70.9 | 0.048<br>28.6 | ~ 0<br>0.05 | 0.9987 | 3789 |
| *double exp.* | 0.31<br>68.18 | 0.045<br>31.82 | / | 0.9971 | 6046 |
| $S1_1$, N= 345<br>[a] *triple exp.* | 0.25<br>66.95 | 0.05<br>32.90 | ~ 4.930e-032<br>0.15 | 0.9984 | 4037 |
| [a] *triple exp.*<br>*weight 1/t* | n.a. | n.a. | n.a. | n.a. | n.a. |
| *double exp.* | 0.25<br>67.98 | 0.048<br>32.02 | / | 0.9983 | 4150 |
| $S1_2$, N=1964<br>*triple exp.* | 0.37<br>35.0 | 0.071<br>59.5 | 0.019<br>5.6 | 0.9998 | 10436 |
| *triple exp.*<br>*weight 1/t* | 1.1<br>(1.0 to 1.2)[b]<br>11.4<br>(10.5 to 12.3)[b] | 0.20<br>(0.19 to 0.21)[b]<br>43.6 | 0.05<br>0.049 to 0.051[b]<br>45<br>(44 to 47)[b] | 0.9998 | 9810 |
| *double exp.* | 0.24<br>49.89 | 0.050<br>50.11 | / | 0.9989 | 18320 |

[a] fitting did not converge properly or was ambiguous

[b] 95 % CI

Notably, however, due to the cumulative distribution plots, the data are dominated by events at short time and with increasing interdependence of the data with increasing time. In order to investigate if this fact erroneously eliminates a dominating fast phase in the non-linear regression fit, we also fitted the data with increased weight (proportional to 1/time) for short times. Strikingly, however, for majority of these cases fitting was ambiguous and complete confidence intervals could not be calculated (Table S1). An exception was the S1 ATPase2 where fitting was successful, showing a fast phase with $k_{\text{fast}}$ = ~1.1 $s^{-1}$ (~15 %). It seems that when enough dwell times are collected (in this case almost 2000) and fitting is weighted (1/time), a faster phase can be discerned. However, importantly, the latter phase is of low amplitude, far from the dominating role in previous work (and our experiments in the IVMA solution). Furthermore, the rate constant is less than half of that found previously and not more than about twice that of our unexplained phase (0.2-0.5 $s^{-1}$). Neither was the fast phase resurrected by limiting the fitting to times <10 s without weighting.

We also used the optimized assay conditions to study single molecule HMM-actin-activated ATPase (Fig. 1g-i; Fig S8). For both dwell-time data-sets triple exponential functions gave good fits with the fastest rate constant attributed to actin-activated ATP turnover.

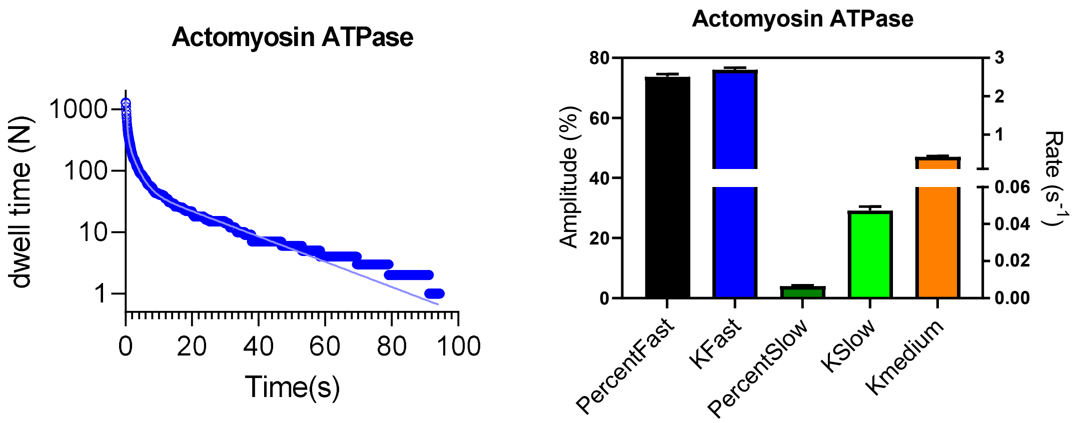


**Fig. S8**. **Reproducibility of optimized single molecule actomyosin ATPase assay.** *Left*: Cumulative frequency distribution of Alexa-nucleotide dwell-time events of actomyosin ATPase activity under optimized conditions. The distributions are fitted by a triple exponential function (solid line). *Right:* Amplitudes and rate constants obtained from the fitting of data on the left. Note that the values are similar to the actomyosin data in Fig 1 (main text) but they refer to other experiments. Columns represent means and error estimates refer to 95 % confidence intervals. Temperature: 20 °C

*Effects of Alexa-ADP*

One basis for the unexplained phase (0.2-0.5 $s^{-1}$) that may be considered is that it is attributable to rebinding of Alexa-ADP. However, if that had been the case its amplitude would increase over the period of observation (15 min) or new very fast phases would be expected to emerge. To test the possibility of Alexa-ADP accumulation we subdivided the 15 min traces into three equal parts (beginning, middle, end) and analyzed them separately. As can be seen from Fig. S11 there was no clear trend to suggest increased amplitude or rate of the fast, unexplained phase over time that might suggest Alexa-ADP accumulation.

To further address the possible contribution of Alexa-ADP binding we created "Alexa-ADP" conditions by pre-incubating HMM with AlexaATP in wash buffer for 1-2 h at room temperature. The reaction mixture was then diluted in TIRF buffer and dwell time events were directly observed. Analysis of extracted traces showed dwell-time distributions with hugely different appearance than in studies of basal HMM ATPase. Particularly, there was an extra dominating fast phase (~2$s^{-1}$) with about 20-fold faster rate than basal HMM ATPase and more than 4-fold faster rate than the unexplained phase in HMM ATPAse (Fig. S9). This further substantiates our conclusion that the unexplained phase observed in basal HMM ATPase is not attributable to (re)binding of Alexa-ADP.

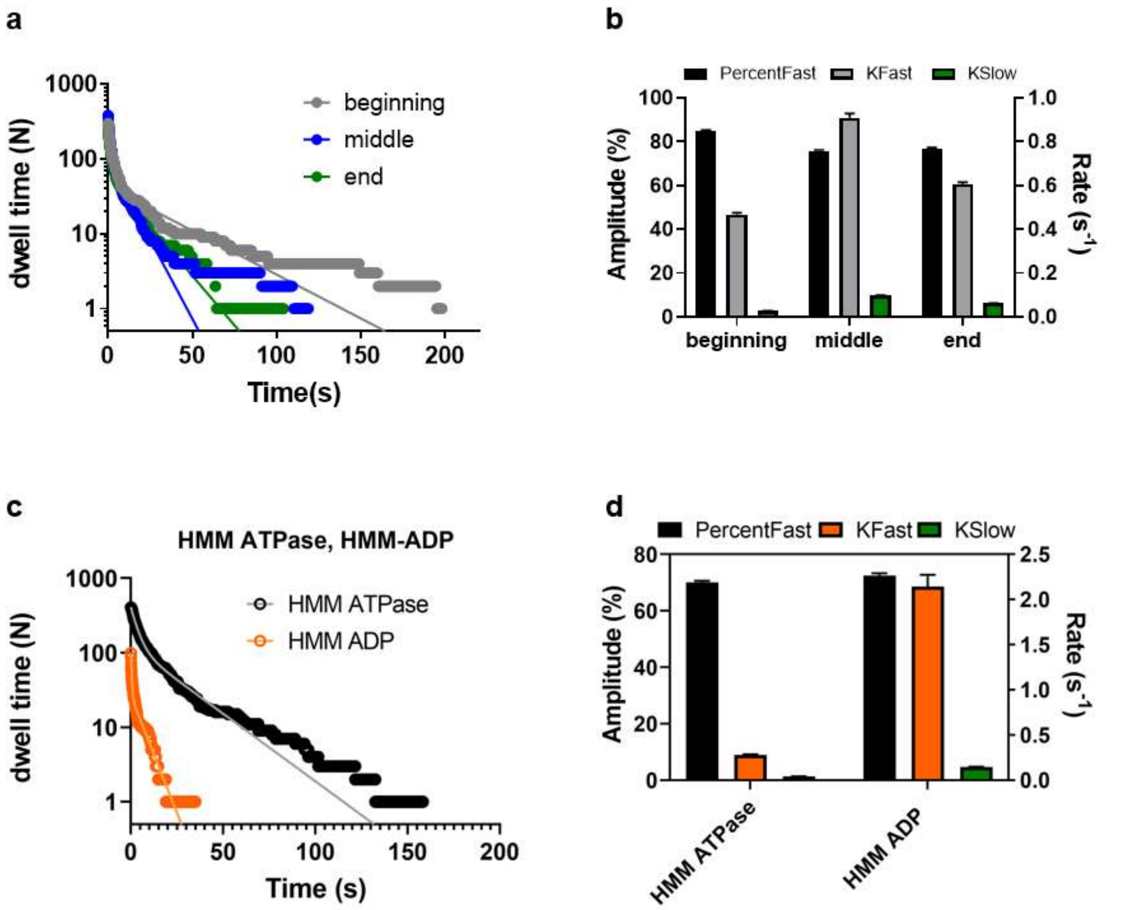


**Fig. S9. Single molecule HMM basal ATPase: role of Alexa-ADP. a**: Cumulative frequency distribution of Alexa647-nucleotide dwell-time events comparing beginning, middle and end part of the 15 min traces. The distributions were well fitted with double exponential functions (solid lines). **b**: Amplitudes and rate constants obtained from the fitting of data to the left. **c**: Cumulative frequency distribution of Alexa647-nucleotide dwell-time events comparing regular HMM ATPase vs events distribution under "ADP conditions" (see text). **d:** Amplitudes and rate constants obtained from the fitting of data on the left. Columns represent means and error estimates refer to 95 % confidence intervals. Temperature: 23 °C.
